# Nonlinear mixed-effect models and tailored parametrization schemes enables integration of single cell and bulk data

**DOI:** 10.64898/2026.04.06.716803

**Authors:** Dantong Wang, Fabian Fröhlich, Paul Stapor, Yannik Schälte, Manuel Huth, Roland Eils, Stefan M. Kallenberger, Jan Hasenauer

## Abstract

Experimental methods for characterizing single cells and cell populations have improved tremendously over the past decades. This progress has enabled the development of quantitative, mechanistic models for cellular processes based on either single cell or bulk data. However, coherent statistical frameworks for the model-based integration of different data types at the single-cell and population levels are still missing.

In this work, we present a mathematical modeling approach for integrating single-cell time-lapse, single-cell snapshot, single-cell time-to-event and population-average data. Utilizing a formulation based on nonlinear mixed-effect modeling, we enable the description of multiple data types, with and without single-cell resolution, and we propose a tailored parameter estimation method. Furthermore, we propose a tailored parameter estimation scheme that facilitates the assessment of underlying process parameters.

Our study demonstrates that the proposed approach can reliably integrate diverse data types, thereby improving parameter identifiability and prediction accuracy. Applying this framework of extrinsic apoptosis reveals that simultaneously considering multiple data types can be essential, particularly when experimental constraints limit data availability. The proposed approach is broadly applicable and may significantly advance our understanding of complex biological processes.

## Introduction

Mechanistic mathematical models are widely employed in the study of biological processes. These models can not only describe key aspects of individual cells and cell populations, but also provide novel insights at both levels. While models for individual cells are typically informed by single-cell time-lapse data (Karlsson et al., 2015; Fröhlich et al., 2018b), models for cell populations rely on single-cell snapshot (Hasenauer et al., 2011; Neuert et al., 2013; Waldherr, 2018) or population-average data (Fröhlich et al., 2018a; Adlung et al., 2021; Gong and Sobie, 2018).

The different types of experimental data can be acquired using a broad spectrum of techniques. For instance, single-cell time-lapse and time-to-event data – which provide a time point when a specific event takes place (e.g., cell division or death) – can be captured by time-lapse (fluorescence) microscopy or time-resolved sequencing methods (Qiu et al., 2020). Single-cell snapshot data can be obtained using techniques such as flow and mass cytometry (Davey and Kell, 1996; Bodenmiller et al., 2012), single-cell sequencing (Frei et al., 2016), or single-cell proteomics (Palit et al., 2018). Population-average data, on the other hand, are accessible through methods such as bulk sequencing (Li and Wang, 2021), microarray analysis (Daran-Lapujade et al., 2008), and immunoblotting (Kurien and Scofield, 2006). Whereas snapshot and population-average data provide statistics for large populations of cells but do not offer time-courses for individual cells, time-lapse experiments track single cells over time while typically involve fewer cells.

Although these experimental techniques vary in terms of throughput and content, each can provide valuable insights, and some information might only be accessible with one particular method. As a result, it is desirable to integrate information from different techniques (Xiao et al., 2024; Jolasun et al., 2025). Accordingly, researchers have already utilized single-cell time-lapse data under various experimental conditions (Almquist et al., 2015a; Karlsson et al., 2015; Fröhlich et al., 2018b), combined population-average and single-cell snapshot data (Adlung et al., 2021), and even integrated multiple data types (Kallenberger et al., 2014) to estimate the parameters of mechanistic models. These efforts have provided novel biological insight, e.g., regarding the sources of cell-to-cell variability (Almquist et al., 2015a; Karlsson et al., 2015), the structure of cellular processes (Fröhlich et al., 2018b), and cellular decision-making (Kallenberger et al., 2014; Adlung et al., 2021). However, to date, it remains unclear how all of the aforementioned data types, namely single-cell time-lapse data (including time-to-event data), single-cell snapshot data, and population-average data, can be integrated in a statistically coherent manner.

For model-based analysis of cellular processes, multiple modeling frameworks are in use. Individual cells are often described using ordinary differential equations (ODEs), stochastic differential equation models, or Markov jump processes (Wilkinson, 2009). Cell populations are frequently described using partial differential equation (PDE) models (Waldherr, 2018), stochastic branching processes (Loos et al., 2015) or (non-linear) mixed-effect models (MEMs) (Karlsson et al., 2015; Llamosi et al., 2016; Fröhlich et al., 2018b). Nonlinear MEMs are among the most widely used frameworks for the study of heterogeneous cell populations, probably because (1) MEMs allow statistically coherent analysis of repeated individual-level observations (Pinheiro, 1994) and (2) computational methods are readliy available. Laplace approximation (Tierney and Kadane, 1986), first-order conditional estimation (FOCE) (Beal and Sheiner, 1992) and stochastic approximation expectation maximization (SAEM) (Kuhn and Lavielle, 2005) support the parameterisation of MEMs based on longitudinal data, as repeatedly demonstrated (Karlsson et al., 2015; Llamosi et al., 2016; Fröhlich et al., 2018b). Nevertheless, the methods are computationally demanding if a large number of individual cells must to be considered, as is the case for single-cell snapshot data, and are not directly applicable to population-average data.

In this study, we propose a framework based on MEMs that enables the integration of diverse data sources, including single-cell time-lapse, single-cell time-to-event, single-cell snapshot data, and population-average data. To this end, we formulate a joint likelihood function that captures the information contained in these data types. We introduce efficient methods to evaluate or at least approximate the individual components of the likelihood function, building on established approaches in the MEM field (e.g., FOCE for single-cell time-lapse data) and developing new approaches (e.g., sigma point (SP) methods (Julier et al., 1995; Menegaz et al., 2011; Lerner, 2002) and Dirac mixture distributions (Gilitschenski and Hanebeck, 2013) for single-cell snapshot and average data. Subsequently, an optimization and uncertainty analysis approach is then introduced to efficiently estimate the unknown parameters of the integrated model and to quantify parameter uncertainties.

To demonstrate the effectiveness of our proposed method, we apply it using synthetic and real-world datasets. In these evaluations, we compare the performance of our integrated data approach with that of methods restricted to using only subsets of the available datasets. The results consistently show that our proposed framework outperforms alternative approaches, leading to more accurate characterization of cell-to-cell variability and improved analysis of sources of cellular heterogeneity.

## Results

### Computational framework for integrating single-cell and population data in mixed effect models

To facilitate the study of cellular processes in heterogeneous populations, we developed a coherent framework for the integration of all commonly available data types for cell populations (Fig. 1a). This framework builds on non-linear MEMs and describes the dynamics of a cell population based on (i) a parametric model for individual cells and (ii) a model for the parameter distribution across the cell population.

**Figure 1:**
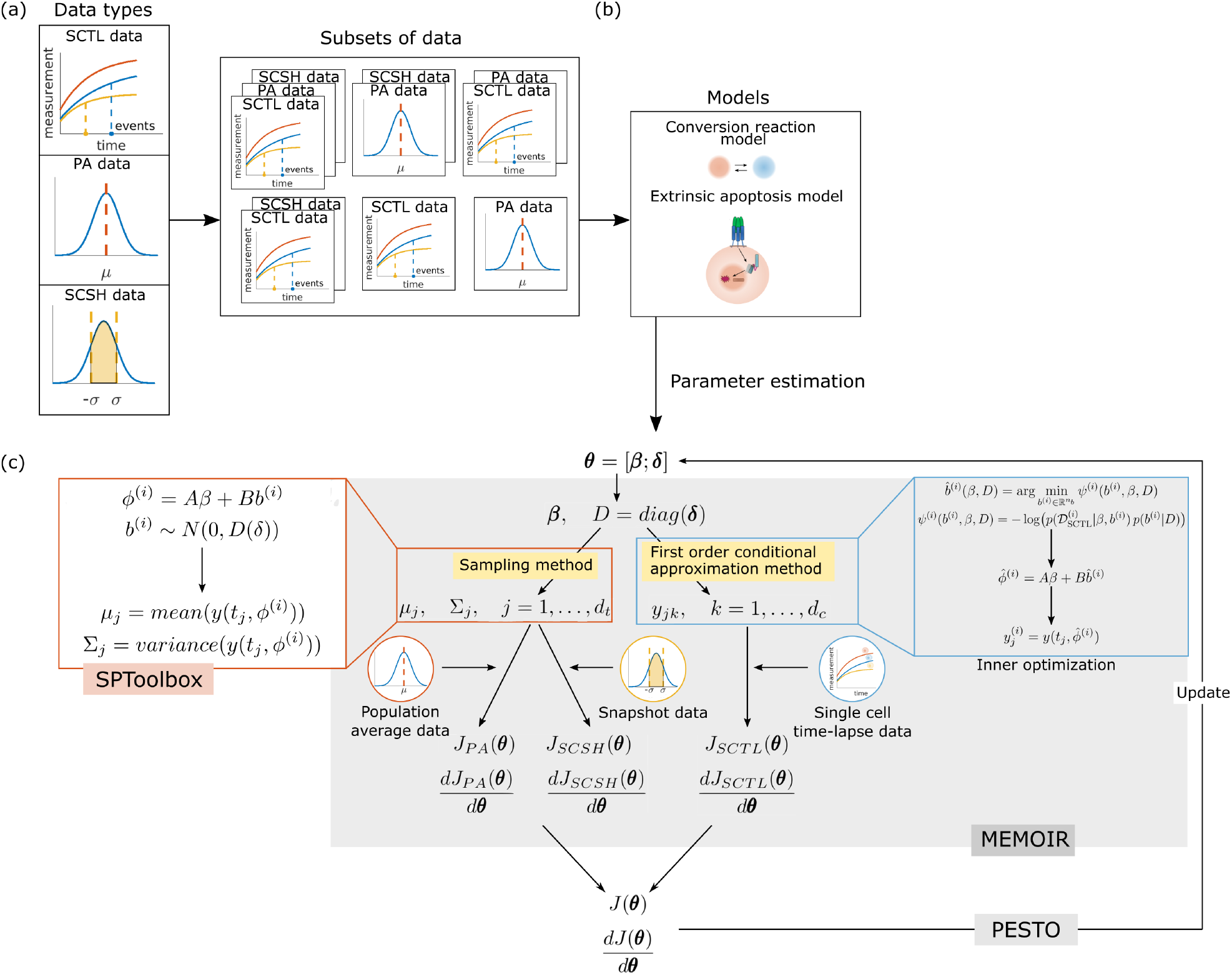
Parameter estimation and comparison of integrating different subsets of data types. (a) Three types are used in this study. Subsets of data are used for parameter optimization separately. (b) Two models are used to access the effects of different data combinations, the conversion reaction model and the extrinsic apoptosis model. The two icons are used in this paper to illustrate which model is used for analysis in the specific figures or sub-figures. (c) Illustration of the optimization routine. The MATLAB toolbox MEMOIR is used to integrate the negative log-likelihood values and the gradients computed for three data types. The negative log-likelihood value of single-cell time-lapse data is computed using the toolbox MEMOIR using the Laplace approximation method. Monte Carlo sampling is used for the population-average and single-cell snapshot data, which is implemented in the MATLAB toolbox SPtoolbox. We used the PESTO toolbox for gradient-based multi-start local optimization.

(i) The **parametric model for individual cells** can, in principle, be any function that maps parameters of individual cells to observables (Fig. 1b). In this study, we focus on mechanistic models of cellular processes based on ODEs,

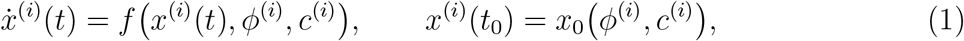

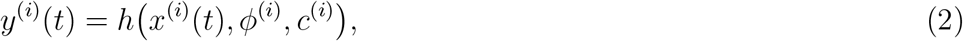

with state *x*^(*i*)^(*t*), observable *y*^(*i*)^(*t*) for cell *i* with parameters *ϕ*^(*i*)^ at time *t*, and covariates *c*^(*i*)^. The time-dependence of the state is described by the vector field *f* — which can be the product of a stoichiometric matrix and flux vector — and the observation process by the observable mapping *h*. The parameters *ϕ*^(*i*)^ are reaction rate constants, initial values or their logarithms. The measurements for individual cells, e.g., in the context of single-cell time-lapse and snapshot measurements, are a noise-corrupted version of the observables at time points *t*_*k*_:

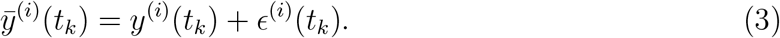

The measurement noise *ϵ*^(*i*)^(*t*_*k*_) might follow, e.g., a normal, log-normal or Laplace distribution, and be correlated. The parameters of the noise model are denoted by Σ. In this case, the parametric model for the relation of cell-specific parameters *ϕ*^(*i*)^ and observations *y*^(*i*)^ is the noise-corrupted evaluation of the ODE solution (operator) for a given observable mapping and time points (see *STAR Methods*).

(ii) The **model for the parameter distribution across the cell population** can be any parametric probability density function (Fig. 1c). For the study of cell populations, normal distributions are frequently used (Llamosi et al., 2016; Fröhlich et al., 2018b). In this case, the parameters of individual cells are determined by fixed effects *β* and random effects *b*^(*i*)^,

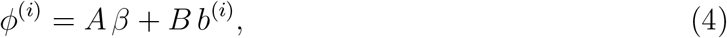

with design matrices *A* and *B*, and the random effect is a realization of a multivariate normal distribution with zero mean and covariance matrix *D, b*^(*i*)^ ∼ 𝒩(0, *D*). Alternatively, distributions with heavier tails or multiple modes can be used (Hasenauer et al., 2011). Symmetry and positive definiteness of the covariance matrix *D* can be achieved through parameterization using a vector *δ* (Pinheiro and Bates, 1995).

A MEM allows for the simulation of a cell population with single-cell resolution as well as the evaluation of population statistics, such as population-averages, based on the parameters *θ* = (*β, δ*, Σ). Hence, while previous publications focused on single-cell time-lapse data, applying MEM also enables the assessment of other data types: here, we formulated the joint likelihood functions for single-cell time-lapse and time-to-event (SCTL) data, single-cell snapshot (SCSH) data, and population-average (PA) data. The datasets are denoted by 𝒟_*SCTL*_, 𝒟_*SCSH*_ and 𝒟_*PA*_ and their collection by 𝒟. Assuming that the individual datasets are independent, meaning that different randomly sampled subpopulations are used for the individual experiments, the joint likelihood function is the product of the likelihood functions for the individual datasets

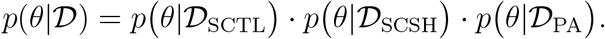

We compute the individual likelihood functions as follows:

#### Single-cell time-lapse (SCTL) data

consist of time series for individual cells. Assuming independence of the single-cell time series — which cannot always be ensured — the likelihood for observing SCTL data is the product of the likelihoods of observing the individual time series. As the random effects for a specific cell are unknown, the likelihood is given by the marginalization of the likelihood for observing the individual time series for a given single-cell parameter *ϕ*^(*i*)^ with respect to the population-level parameter distribution. We approximated these integrals using the Laplace approximation (Tierney and Kadane, 1986), an established method for nonlinear MEMs. To ensure numerical robustness, we employed sensitivity equations for optimization and approximation.

A special case of SCTL data is time-to-event data. Here, events are associated with the dynamics of individual cells through trigger functions (e.g., crossing a threshold). Conceptually, these events can be treated like other single-cell observations; however, in this case, the observed quantity is the time at which the event occurs. As in other quantities, the observed time point may be subject to measurement noise. For the computation of sensitivities, we employ the implicit function theorem (Fröhlich et al., 2016).

#### Single-cell snapshot (SCSH) data

provide time series of population snapshots. Each snapshot at time point *t*_*k*_ contains measurements from a large number of cells, often in the thousands. Under the assumption of independence, one could, in principle, use the same likelihood function formulation as for SCTL data. However, the existing methods for likelihood evaluation or approximation are (i) computationally demanding when cell numbers are large and (ii) unreliable if the information per cell is limited (e.g., when parameters remain non-identifiable). Therefore, in this study, we do not directly use the single-cell data but rather the statistical moments of the data — namely, the time-dependent population mean and variance. The likelihood function is then the conditional probability of observing the empirical estimates of these statistical moments of the SCSH data given the MEM simulation. To compute these statistical moments, we use Monte Carlo sampling.

#### Population-average (PA) data

provide a time series for the mean of the observable. This mean is closely related but not necessarily identical to the mean of the SCSH data. Indeed, the two differ if the cell-level measurement noise has a non-zero mean, which is already the case for multiplicative, log-normally distributed measurement noise. For the SCSH and the PA data, the noise level in the respective likelihood formulations is not based on the uncertainty of the empirical moment estimators, as this appeared overly optimistic. Instead, we use the standard deviation resulting from differences between independent experiments. A mathematically concise formulation is provided in the section *STAR Methods*.

The individual likelihood formulations and the expression for the joint likelihood are a central element of our framework for integrating heterogeneous data types. They provide a systematic approach for evaluating parameterized models and for inferring unknown model parameters, including the fixed effect vector and the covariance matrix. For parameter inference, we employ a gradient-based multi-start local optimization strategy followed by uncertainty analysis. All gradients are computed using forward sensitivity equations, ensuring numerical robustness. Gradient calculation relies on known formulations for SCTL data (Almquist et al., 2015b) and newly derived expressions for the remaining data types. Differentiability of Monte Carlo sampling results is achieved by drawing samples from a multivariate standard normal distribution and transforming them in each iteration with the square root of the covariance matrix. Gradient-based optimization is initialized at randomly sampled starting points and performed using a trust-region reflective method.

### Pathway models and datasets for the assessment of the computational framework

To illustrate the properties of our approach and the effects of different data types in both simple and complex models, we consider two models and corresponding datasets:

First, we consider a **model for a reversible conversion reaction**, *x*_1_ ⇌ *x*_2_. We create three artificial datasets, one for each of the aforementioned data types. Following common real-world setups, we assume that the PA and SCSH data provide absolute measurements (e.g., by using a calibration curve), whereas SCTL data provide only relative measurements. Accordingly, for the SCTL data, we introduce an unknown, experiment-specific scaling factor *ϕ*_3_. The single-cell time-to-event data provide the time point 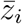 at which the concentration *x*_1_(*t*) reaches the value *x*_1_(0) · *ϕ*_4_. For details regarding the observable and noise model, as well as the parameter values used for data generation, we refer to the *STAR Methods*.

Second, we consider a **model of extrinsic apoptosis**, which links CD95L concentrations to apoptosis kinetics and fractional killing in a heterogeneous cell population. The model describes the binding of CD95L to the cell death receptor CD95, leading to the formation of active death receptors (Fig. 2) and the recruitment of the adaptor protein Fas-associated death domain protein (FADD). This recruitment results in the formation of death-inducing signaling complexes (DISCs) (see Table S1 for model equations). Within DISCs, dimerized procaspase-8 (p55) is cleaved to the fragments p43 and p30 (Hoffmann et al., 2009). Subsequently, p43 and p30 are further cleaved to p18, which then dissociates from the DISC and enters the cytosol. The catalytically active forms of caspase-8 (p43 and p18) can cleave BID (BH3 interacting-domain death agonist). Truncated BID (tBID) then accumulates in mitochondria. If a certain threshold concentration of tBID is exceeded, mitochondrial outer membrane permeabilization (MOMP) ensues, resulting in apoptosis.

**Figure 2:**
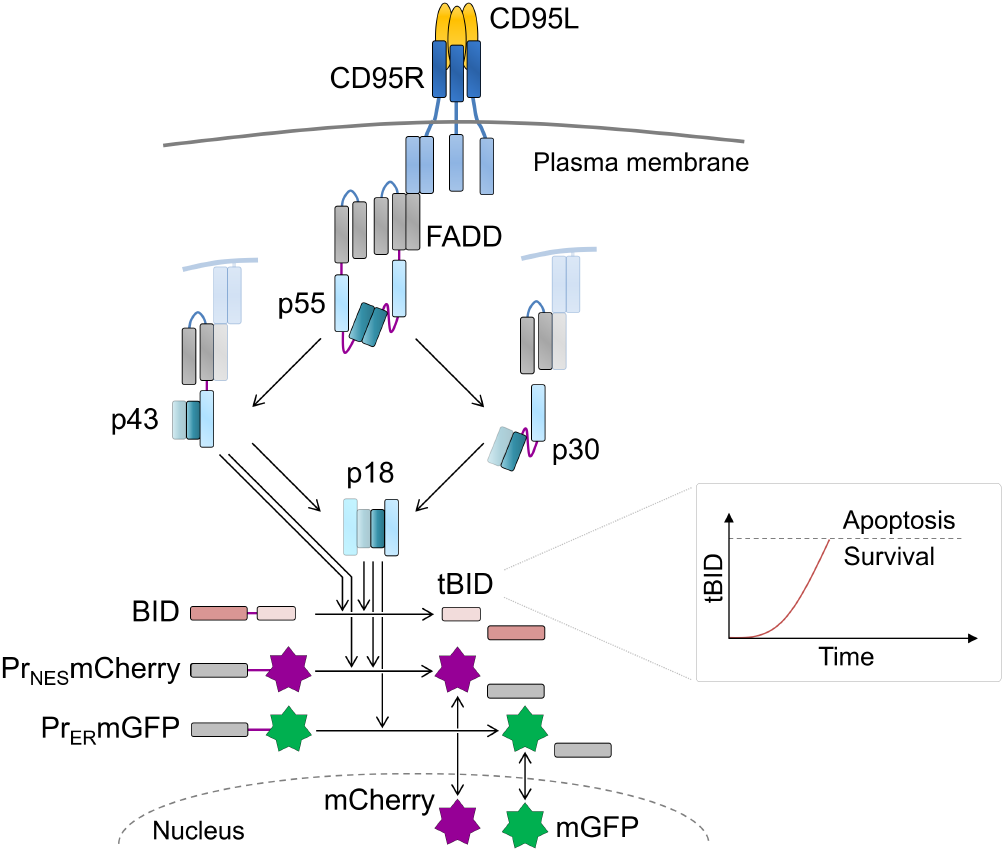
Model of extrinsic apoptosis. Binding of CD95L to CD95R results in assembly of complexes with FADD and dimers of procaspase-8 (p55). Cutting at two cleavage sites, drawn in purple, results in p43 or p30, and finally in p18. Membrane-bound p43 and cytosolic p18 are catalytically active and can cleave BID to tBID as well as the cytosolic probe *Pr*_*NES*_*mCherry*. The ER-bound probe *PR*_*ER*_*mGFP* can only be cleaved by p18. After cleavage of probes, mCherry and mGFP can enter the nucleus, which is detected by confocal microscopy. Inlay: in case, a certain threshold of tBID is exceeded, a cell undergoes apoptosis.

For this model of extrinsic apoptosis, we consider experimental datasets for wild-type HeLa cells and stably CD95-overexpressing HeLa cells (CD95-HeLa) (Kallenberger et al., 2014). Using fluorescent cleavage probes, cleavage activities of p18 and p43 were quantified in single cells. A cytoplasmic probe, *Pr*_*NES*_*mCherry*, indicated the activity of p18 and p43, whereas an ER-bound probe, *PR*_*ER*_*mGFP*, indicated only the activity of p18 (see *STAR Methods* for details). The two cell lines were treated with CD95L at four different concentrations, yielding distinct apoptosis kinetics. Compared to our previous study, the model fitting approaches were here applied to a larger collection of experimental data. Specifically, for model calibration, 20 additional single-cell trajectories from CD95-HeLa cells were included, resulting in a total of 30 single-cell trajectories per ligand concentration (instead of 10). Due to the accelerated cell death kinetics in these cells, more single-cell trajectories could be recorded and used for model fitting (*n* = 30 per ligand concentration) than for wild-type HeLa cells (*n* = 10 per ligand concentration). Accordingly, the MEM was fitted to a total of 160 single-cell trajectories. In addition, single-cell apoptosis times, quantitative immunoblots, and fluorescence-assisted cell sorting measurements (representing single-cell snapshot data) were used for model fitting. Further details of experimental procedures, data processing, and the definitions of model observables are described in the *STAR Methods*. In-depth descriptions of the experimental methods can be found in Kallenberger et al. (2014). Collectively, 14 datasets, including 8 single-cell time-lapse datasets, 4 population-average datasets, and 2 single-cell snapshot datasets, were combined for model fitting. An overview of all datasets is provided in Table S2.

In the following, we first use these two models to assess the critical components of the computational pipeline and validate our implementation. Subsequently, we demonstrate the benefits of coherent data integration.

### Gradient-based multi-start local optimization facilitates robust likelihood evaluation for single-cell time-lapse data

The optimization of the model parameters requires the evaluation of the individual likelihood contributions. For single-cell time-lapse data, the likelihood is approximated using the Laplace approximation. This integral approximation exploits the most likely value of the random effects, 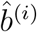, using the empirical Bayes optimization problem (Lindstrom and Bates, 1990). This inner optimization problem must be solved for each single cell in each outer optimization step for the population-level parameters, *β* and *D*. For the outer optimization to converge, the inner optimization method must be highly reliable. In this study, we used gradient-based, multi-start local optimization to solve the inner optimization problem.

To assess the convergence of the inner optimization and the resulting stability of the gradient, we performed 500 local optimization runs for each SCTL dataset. For the model of extrinsic apoptosis—the more challenging application—we found that, for fixed values of *β* and *D*, at least 188 out of the 500 local optimization runs converged (Fig. S1), implying a success rate of 37.6%. A single optimization run is therefore insufficient; however, if 20 starts are performed, the probability that at least one run converges is at least 99.99% (calculated as 1 − (1 − 37.6%)^20^), which is reasonably high. To further increase the convergence rate, we initiated one start at the optimal point found in the previous step of the outer optimization, which further boosted both inner and outer convergence.

### Monte Carlo sampling provides estimates for statistical moments and enable robust likelihood evaluation for single-cell snapshot and population average data

The likelihood of single-cell snapshot and population average data is formulated using the statistical moments of the mixed-effect model, namely the mean and variance. Since the robust evaluation of these quantities is essential, different numerical methods have been developed, each trading off approximation accuracy against computational cost. Among these methods, Monte Carlo sampling (MC) provides a flexible framework by repeatedly drawing samples from the random parameter distribution, whereas the sigma point (SP) method employs a fixed set of sample points to approximate the mean and variance. Given the limitations of available assessments [e.g., Wang et al. (2019)], we implemented both approaches in the computational pipeline and assessed how accurate and efficient they are for our models.

To quantify the differences in accuracy, we used a high-fidelity Monte Carlo simulation with 100,000 samples to approximate the true population mean and variance. For each tested method (SP or MC with fewer samples), we then computed the relative error of the approximated mean (or variance).

Our assessment revealed that the SP method, which uses a fixed number of samples, struggles to capture the full density of the observable distribution (Figs. 3a,b). Indeed, tails are mostly underrepresented. This results in approximation errors which can be rather severe (Figs. 3c,d).

**Figure 3:**
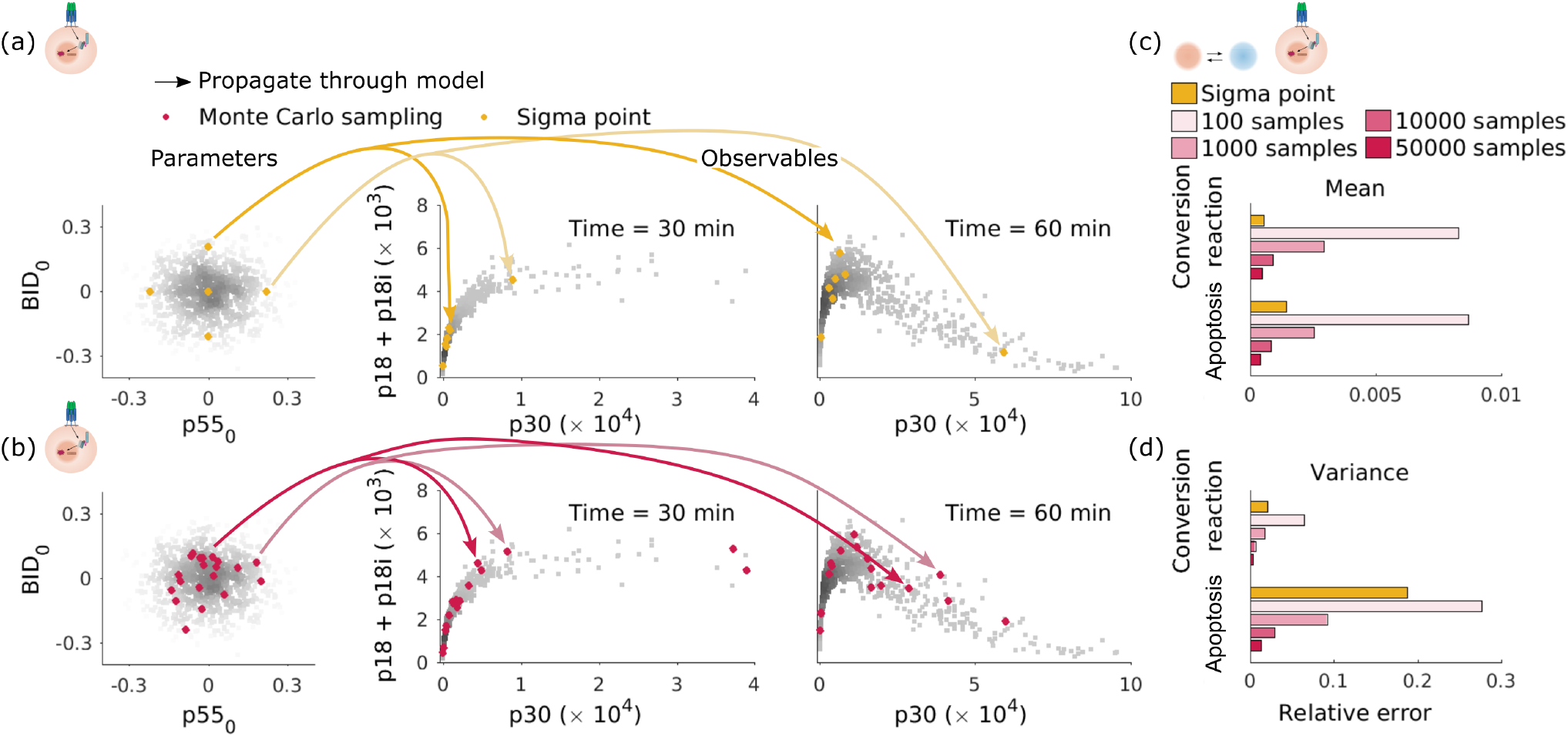
Comparison of sigma point and Monte Carlo sampling method. (a) Sigma point (yellow) samples of parameters propagate through the model and output values at different time points. We use the two parameters *BID*_0_ and *p*55 as an example. The five sigma point samples are propagated through the model and the corresponding observables are plotted. The correlation of two observables is shown using 1,000 Monte Carlo samples (grey) at two time points, 30min, and 60min for comparison. (b) Monte Carlo samples (red) of parameters propagate through the model output values at different time points. To avoid overlapping each other, here only 20 Monte Carlo samples are shown to compare with the distribution predicted by sigma point methods. The relative error of the mean (c) and variance (d) was predicted using sigma point and Monte Carlo sampling methods with different sample sizes.

For the model of the conversion reaction—comparatively simple and linear—, the accuracy of the SP method was comparable to MC sampling with 10,000 draws. The SP method achieved for the mean a relative error of 0.05%, while the SP method achieved a relative error of 0.09%. For the variance, we observed a relative error of 2% versus 0.59% in favor of the MC method. For the model of extrinsic apoptosis, the SP method yielded substantially larger errors, especially for the variance (18.7% for the SP method versus 2.9% for the MC method), even compared to MC with 10,000 samples. We further examined whether increasing the MC sample size beyond 10,000 improves accuracy. Thus, for simpler models, the SP method can be suitable, but for complex models MC sampling is required.

### Gradient computed by forward sensitivity method is accurate and efficient

For efficient and robust evaluation of the likelihood function is of critical importance. Yet, various studies showed that for high-dimensional estimation problems, also robust gradient calculation is required (Raue et al., 2013; Fröhlich et al., 2018a; Villaverde et al., 2019). In this study, we provide a comprehensive mathematical formulation of the likelihood function gradients based on sensitivities equations and an implementation of this approach in the afore-describe computational framework. Here, we conducted a thorough evaluation of this implementation by comparing gradients calculated via forward sensitivities—the method we use during optimization—to those obtained via forward finite differences, and evaluated the respective computation times. We performed this evaluation using 20 multi-starts for the inner optimization problem and 10,000 Monte Carlo samples for moment approximation. The Monte Carlo samples are not redrawn between optimizer step but just transformed according to the covariance matrix *D*. The same setup was chosen in the following parts of the manuscript.

For the conversion reaction model, our evaluation revealed an excellent agreement between the gradient values computed using forward sensitivities and forward finite differences for the log-likelihood functions of SCTL, SCSH, and PA data (Fig. 4a). As the accuracy of forward finite differences depends on the step size, the level of agreement between the two methods does as well (Fig. 4b). For SCSH data, the best agreement was achieved with a step size of 10^−7^, whereas for SCTL data, the best agreement was obtained with a step size of 10^−3^. Hence, the step size yielding the minimal approximation error of finite difference-based gradient calculation depends on the data type, highlighting the advantage of forward sensitivity analysis, which leverages adaptive numerical solvers. In addition, the computation time for forward sensitivities was generally lower than for finite differences (Fig. 4c). Indeed, only for the already very fast evaluation of PA data, we observed a slight overhead. As the overall computational cost was dominated by the evaluation of the likelihood of the SCTL data, gradient calculation through forward sensitivities was overall 5-times more efficient than through finite differences.

**Figure 4:**
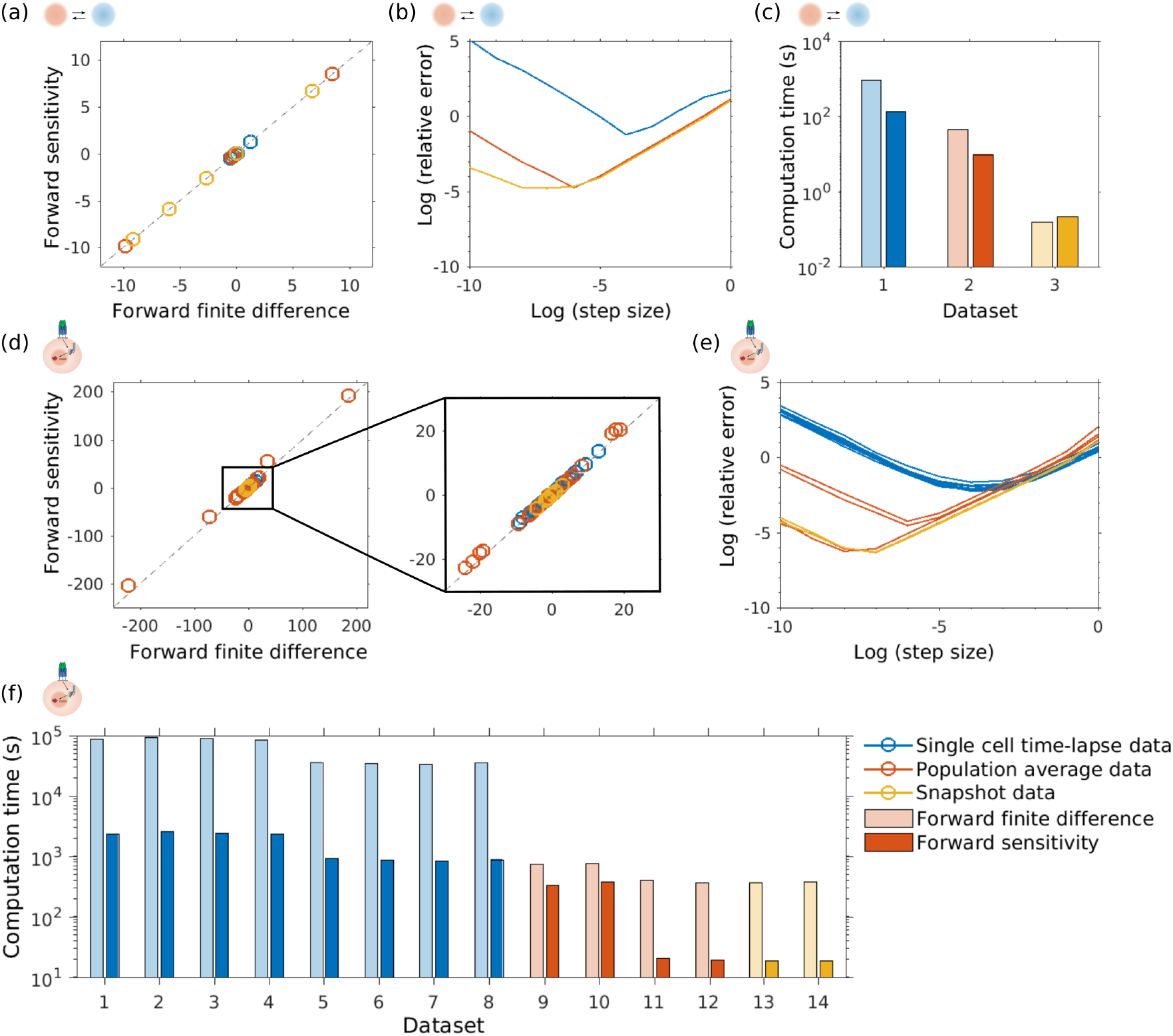
Comparison of computing gradient values using forward sensitivity and forward finite difference method. (a) Comparison of gradient values computed by forward sensitivity and forward finite difference method with the step size 10^−4^ for the conversion reaction model. Each point represents the gradient of one negative log-likelihood function with respect to one parameter. (b) Relative error of gradient computed by forward finite difference method for different step sizes, the ground truth is computed using forward sensitivity method. (c) Computation time of the two methods for 3 datasets with different data types. (d) Comparison of gradient values computed by forward sensitivity and forward finite difference method with the step size 10^−2^ for the extrinsic apoptosis model. Each point represents the gradient of one negative log-likelihood function with respect to one parameter. (e) Relative error of gradient computed by forward finite difference method with different step sizes. The individual lines indicate different datasets of a specific data type. (f) Computation time of the two methods for all 14 datasets.

For the model of extrinsic apoptosis, we observed similar results to those from the conversion reaction model (Fig. 4d-e). Nonetheless, there are larger differences between the gradient values computed through forward sensitivities and forward finite differences (Fig. 4d). Moreover, the difference in computation times is more pronounced, with forward sensitivities being nearly two orders of magnitude faster (Fig. 4e). These more pronounced discrepancies likely stem from the increased complexity of the numerical simulations.

Overall, our analysis demonstrates that gradient computation via the forward sensitivity method is both accurate and efficient. Furthermore, it remains preferable because it does not require the specification of a step size, which can vary depending on the data type.

### Joint modeling of different data types improves the predictive power of population models

Different data types may vary in their informativeness for model predictions. To study the benefits of integrating different data types, we assessed the predictive power of models trained with different combinations of datasets: (1) all datasets, (2) single-cell time-lapse data left out, (3) population-average data left out, (4) single-cell snapshot data left out, (5) both population-average and snapshot data left out (for conversion reaction model), (6) both single-cell time-lapse and snapshot data left out (for conversion reaction model), (7) wild type cell line left out (for the apoptosis model). The datasets that were disregarded in the training process were used to determine the relative information content of the trained models. We performed the analysis for the model of the conversion reaction as well as the model of extrinsic apoptosis, and refer to the supplement for an assessment of the parameter optimization performance (Fig. S3).

For the model of the conversion reaction, we observed a good agreement between the training data and the model fit for all scenarios (Fig. 5a). This indicates that parameter estimation was functioning properly. Moreover, the predictions of the trained models agreed well with the retained validation data in several scenarios (Fig. 5a, green lines). However, using only population-average data was—for obvious reasons—insufficient to predict the variability observed in single-cell snapshot data (scenario 6). Furthermore, we found that single-cell time-lapse and time-to-event data cannot be predicted from the remaining datasets (scenarios 2 and 6) due to missing scale information. In fact, providing the scaling factor for single-cell time-lapse data and the threshold for time-to-event data improved predictions in these scenarios.

**Figure 5:**
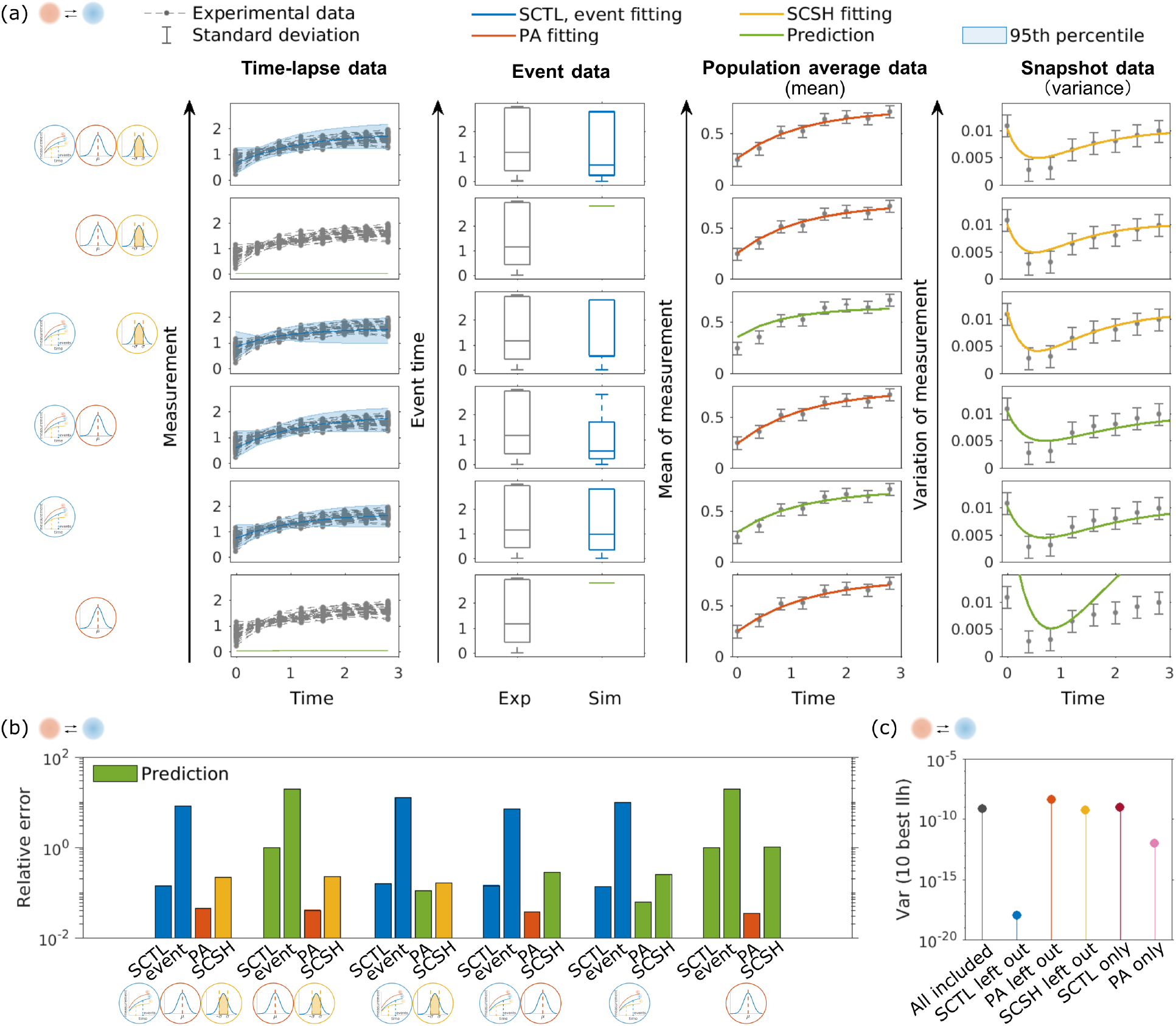
Assessment of the predictive power of the model for the conversion reaction trained using different dataset combination. (a) Fitting the conversion reaction model using different data type combinations. Top to bottom: Different scenarios of including/excluding data in the optimization. Left to right: Measured data (black), fits (data type specific colors), and predictions (green) for time-lapse, event, population-average, and snapshot data. The standard error of the population-average data (second to right) and the snapshot data (right) are shown in black. The interval between the 5th and the 95th percentile of the errors across 1000 realization of single cells are depicted (left) in blue shadow. Datasets not used for training but only for testing are indicated in green. (b) Relative error of fitted (data type specific colors) and predicted (green) datasets compared to the artificial data. (c) Variance of 10 best log-likelihood values of different data combinations.

An in-depth analysis of the relative errors revealed that, as soon as one dataset was excluded, the error between model simulations/predictions and data increased (Fig. 5b). In case of this model, the SCTL dataset was most valuable (with time-to-event data) because fitting to this dataset alone allowed for a reasonable prediction of the remaining data types up to scaling factors. Interestingly, training with single-cell time-lapse and time-to-event data appeared to be particularly challenging, as indicated by the reduced variance of the optimization results when they are excluded.

For the extrinsic apoptosis model, we considered—due to our experiences with the model for a conversion reaction, namely the limited reconstruction accuracy which was achieved when only a single dataset was used—scenarios 1 to 4 and 7. Again, we observed for all scenarios a good agreement of training data and model fit (Fig. 6a). However, also for this real-world dataset, the predictions of the trained models did not agree with the retained validation data (Fig. 6a, green datasets). In scenarios 3 and 5, the excluded data could not be predicted using the remaining data types (Fig. 6a) because some parameters were not shared between models for different data types or cell lines. When the snapshot data were left out, the fitting of the event data became worse. Further, the SCTL dataset from wild-type HeLa cells could not be predicted from the model exclusively calibrated with data for CD95-HeLa, arguably due to cell line-specific parameters for initial protein concentrations. In principle, if information of those parameters were available from other sources, data of the corresponding cell line could be predicted.

**Figure 6:**
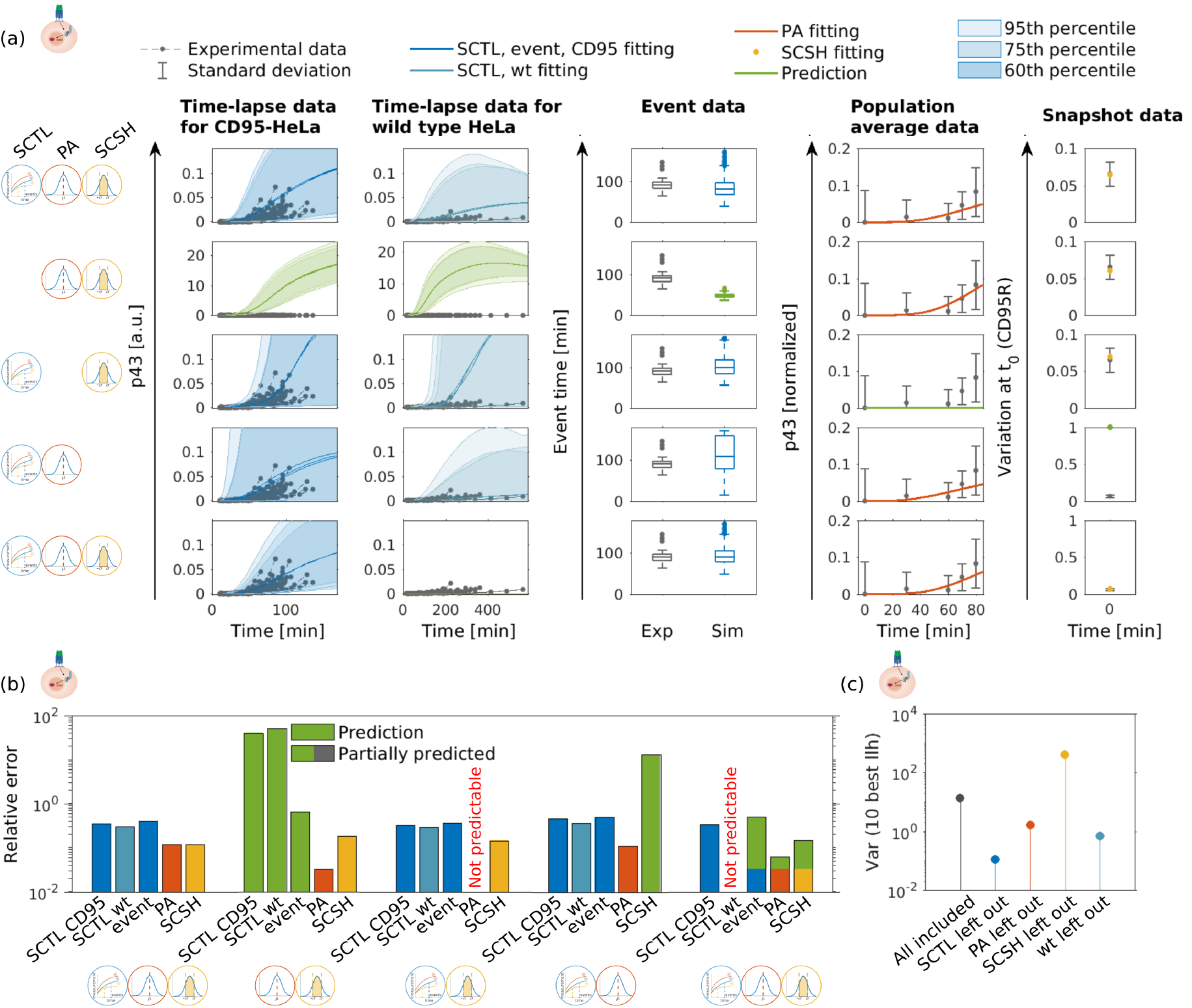
Comparison of different data combinations for extrinsic apoptosis model. (a) Fitting the apoptosis model using different data type combinations. Top to bottom: different scenarios of including/excluding data in the optimization. Left to right: measured data (black), fits (colors indicate data types), and predictions (green) for time-lapse, event, population-average, and snapshot data. The standard error of the population-average data (second right) and the snapshot data (right) are shown in black. The interval between the 20th and 80th, 12.5th and 87.5th, 5th and the 95th percentile of the errors across 1,000 realization of single cells are depicted (left for the CD95-HeLa cell line and second left for the wild type HeLa cell line) in blue shadow. (b) Relative error of fitted (data type specific colors) and predicted (green) datasets compared to the experimental data. The partial prediction refer to the left-out cell type. (c) Variance of 10 best log-likelihood of different data combinations.

The results of the visual inspection (Fig. 6a) are confirmed by the assessment of the relative errors for the different data types (Fig. 6b). Predictions for the retained datasets were either not possible for resulted in large errors. In line with the results for the model of the conversion reaction, the optimizer convergence was best for the case without SCTL data (Fig. 6c). Yet, as these data were required for reliable predictions, disregarding this dataset was not possible. When SCSH data were left out, the convergence performance became worse, which was also consistent with Fig. 6a (scenario 4).

In summary, the analysis of the considered examples demonstrated that the different data types provided complementary information. The joint use of the data improved the predictive performance of the models (Fig. S2). While optimization converged better if merely a subset of the data types was considered, this was not practical and underlined the importance of computational pipelines for data integration.

### Integrating all data types improves parameter identifiability

To investigate how different data combinations affect the parameter identifiability, we assessed parameter uncertainty of the conversion reaction model by computing profile likelihoods (Raue et al., 2013) for all parameters (Fig. 7). When the single-cell time-lapse data were included, all parameters were identifiable (scenarios 1, 3, 4, 5). The parameters estimated using all data types are close to the true parameter values with relative small confidence intervals (scenario 1), compared to the other data type combinations. With only the time-lapse data left out, the confidence interval of parameter ***θ*** = [*θ*_1_, *θ*_2_] was roughly the same as when using all data types (scenario 2), but the event threshold and scaling factor were unidentifiable, which is consistent with the predictions in Fig. 5. Leaving out the snapshot data yielded better parameter identifiability than leaving out the population data (scenarios 3 and 4), indicating that the population-average data contains more information about the parameters. With only time-lapse data included, all parameters were still identifiable, but with much wider confidence intervals (scenario 5).

**Figure 7:**
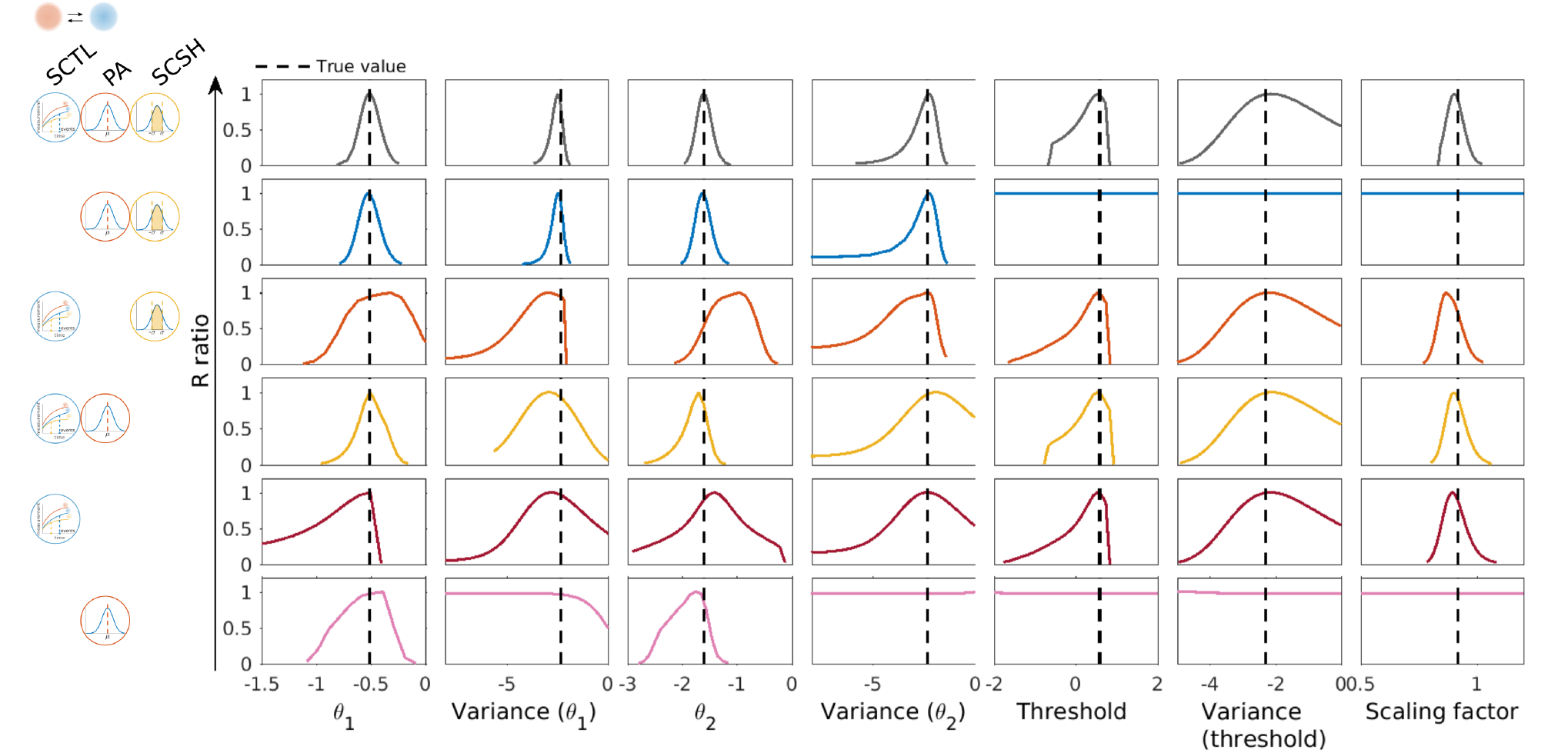
Profile likelihood of all parameters optimized using different data type combinations. Top to bottom: different scenarios of including/excluding data in the optimization. Left to right: model parameters of the forward (*θ*_1_) and backward reaction (*θ*_2_). The true parameter values are indicated by black vertical lines. Curves with the data type specific colors indicate the calculated profiles started at the optimized parameters (usually at the peak of the curves).

Profile calculation would have been prohibitively computationally expensive for the apoptosis model. Instead, we quantified the parameter uncertainty via eigenvalues of the Hessian matrices at the optimal parameter vector, using a local approximation (Fröhlich et al., 2018a). Estimates of confidence intervals can be found in Table S3. In Fig. 8a (right column), the spectrum of the 35 eigenvalues per model was visualized via a kernel density estimator, small eigenvalues indicating higher uncertainties. As before, the uncertainty was the smallest when all data types were included in the optimization process (1st row). When datasets from the wild type cell line were left out, the number of the eigenvalues smaller than 10^−5^ was higher because the cell line-specific parameters could not be optimized at all. However, the median of the eigenvalue spectrum was not substantially decreased because still all data types were used for fitting (2nd row). Similar to the above-mentioned conversion reaction model, leaving out the PA data yielded more eigenvalues numerically close to 0 (*<* 10^−5^) than leaving out the SCSH data (3rd and 4th row). When we left out the time-lapse data, the parameter uncertainty was the largest. In line with Fig. 6c, when we left out the SCSH data, the fluctuation of the median values was the largest implying that the estimated parameter values were largely different between optimization runs.

**Figure 8:**
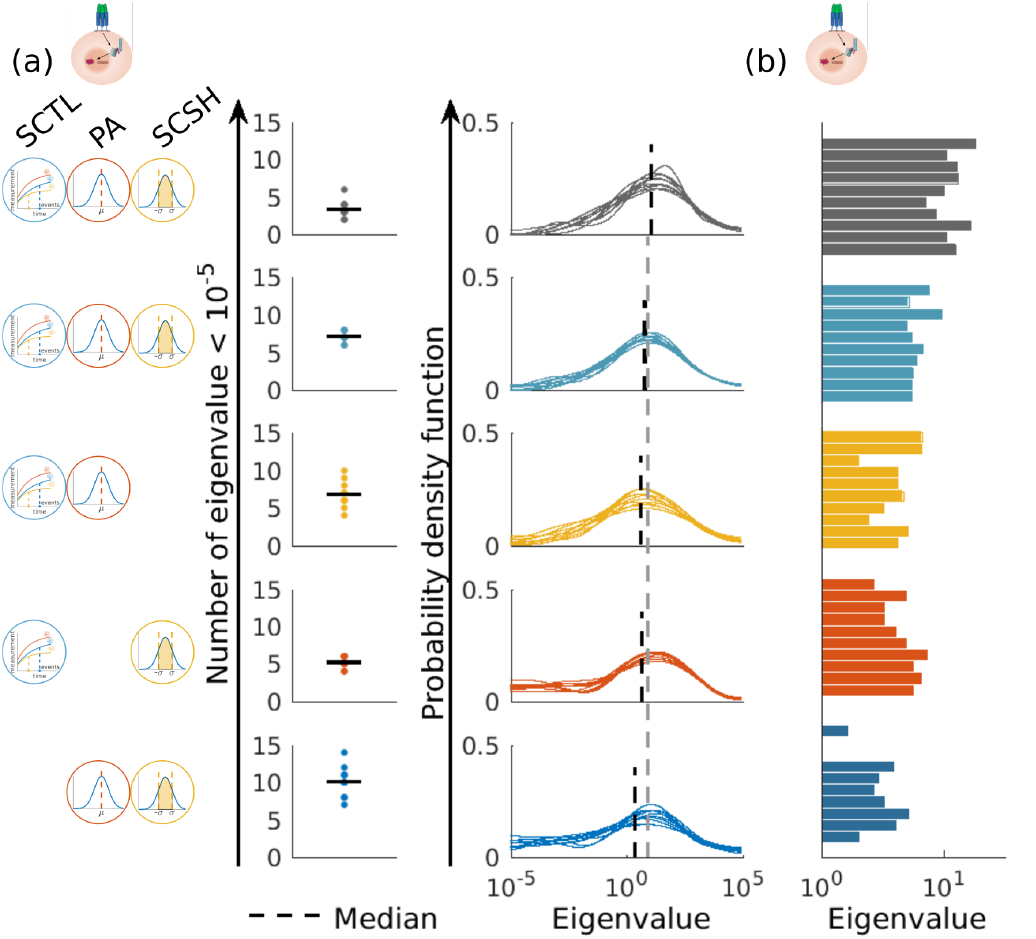
Parameter identifiability for the extrinsic apoptosis model assessed via Hessian eigenvalues. (a) Eigenvalue probability density of the Hessian matrices. Frequency of eigenvalues below 10^−5^ (out of 35 eigenvalues) are plotted on the left. The probability density functions of the best 10 optimization runs for different data type combinations are shown in the right column, restricted to values above 10^−5^, using density plots. The median values of the frequency and the probability density function are depicted as black (dotted and solid) lines. Smaller eigenvalues correspond to larger uncertainties. (b) Median of the eigenvalues of the 10 best runs.

In conclusion, it was necessary to include all three data types to ensure the best parameter identifiability. By comparing different combinations using the two models, we showed that the parameter identifiability was the worst when SCTL data were excluded, while leaving out the PA data yields the second worst identifiability.

## Discussion

Cell-to-cell variability is widely observed in various biological experiments. Non-linear MEMs are a powerful approach to account for this variability in mechanistic models. Depending on the experimental method, different types of data can be measured, giving measurements resolved on the single-cell level, or only population statistics. In this study, we demonstrated that different data types can be integrated into a mixed effects modeling framework. Specifically, we integrated single-cell time-lapse data, population-average data, and snapshot data. To fit parameters, we employed a gradient-based optimization approach. We applied the method to two problems, a simple conversion reaction test model, and a model of extrinsic apoptosis. We demonstrated substantially improved performance in terms of data fits and parameter identifiability when combining all three data types, compared to only a subset. Besides, we also estimated parameters of two cell lines simultaneously in the same model. We showed that it was possible to define both cell line-specific and shared parameters in MEMs, whereas, without knowledge of the cell line specific parameters, data of the corresponding cell line could not be predicted.

A limitation of our approach is that we effectively weight all datasets equally, meaning that if we have more measurements for one data type, it has a higher impact on the result. In our optimization routine, it is possible to estimate parameters shared and independent in different cell lines simultaneously. The same method of defining parameters can also be used for single-cell measurements with subgroups. It is also possible to weight different cell lines (subgroups) differently. An interesting direction for further research would be to investigate methods that choose weights to balance the impact of, e.g., different data types and cell lines (Schaelte and Hasenauer, 2022), however, a fair weighting of heterogeneous data is an open problem. In addition, one might want to penalize deviations from the inferred random-effects distribution from the covariance matrix *D*. Although this was not necessary for the considered application examples, we provide a formulation of a potential penalty term (see *STAR Methods* for details).

For the considered application examples, we assumed that the covariance matrix *D* of the random effects is diagonal, i.e. there is no correlation between random effects, implying the assumption that all initial values are independent in our biological model. Yet, the proposed computational framework and its implementation allow for arbitrary positive definite covariance matrices. This increases the parameter space and the computational complexity, but does not result in a conceptual change.

Our method was applied in two models with different complexity, the simple model of a conversion reaction, and the extrinsic apoptosis model. To ensure the approximation accuracy of the population-average and single-cell snapshot data, we used Monte Carlo sampling method with 10,000 samples instead of SP methods. However, for the simple conversion reaction model, the SP method can potentially decrease the computation time while keeping the approximation accuracy at the same time (Fig. 3). As an alternative, the CMD method (Wang et al., 2019) allows to flexibly increase the approximation accuracy and could conceptually be combined with our data integration approach in the future. Regarding the single-cell time-lapse data, our approach scales linearly with respect to the number of single cells. Nevertheless, the application to higher throughput experiments can be problematic because of the computation time for hundreds of cells. One could investigate methods using, e.g., subsampling or mini-batch approaches (Stapor et al., 2022) to scale to such scenarios. To further accelerate inference and facilitate a more comprehensive uncertainty quantification, also novel neural posterior estimation techniques based on invertiable neural networks or diffusion models could be employ (Radev et al., 2020). These methods have been introduced for standard mixed effect modeling frameworks (Arruda et al., 2024) and could potentially be extended to the more flexible framework presented her.

In summary, we developed an approach to integrate different data types in a mixed-effect modeling framework. On a test and an application problem, we demonstrated improved data fits and parameter identifiability, compared to only using a subset of the available data. Therefore, by including more data types in a mechanistic model, more insight can be gained from different aspects. We anticipate that such approaches will improve our understanding of complex biological pathways by leveraging information from different sources.

## Resource availability

### Lead contact

Further information and requests for resources and reagents should be directed to and will be fulfilled by the lead contact Jan Hasenauer.

### Materials availability

This study did not generate new materials.

### Data and code availability

We implemented the joint log-likelihood and its gradient integrating the three data types in the MATLAB toolbox, building on the MEMOIR toolbox (https://github.com/ICB-DCM/MEMOIR). The Monte Carlo sampling method to approximate the negative log-likelihood for population average and snapshot data is implemented in the MATLAB toolbox SPToolbox (https://github.com/ICB-DCM/SPToolbox), where we also implemented the sigma point method for comparison. To solve the optimization problem, we employed multi-start local optimization via the open-source MATLAB toolbox PESTO (https://github.com/ICB-DCM/PESTO) (Stapor et al., 2018). The local optimization was performed using the MATLAB function `fmincon` with the interior-point method. All codes used for this study are available at https://github.com/ICB-DCM/Extrinsic-apoptosis-Conversion-reaction-model. For profile calculation of the conversion reaction model, we used a method employing iterative optimization along profiles (Raue et al., 2009), also implemented in PESTO.

## Acknowledgement

DW is supported by the China Scholarship Council (201706060200). PS, YS, MH, and JH are supported by the German Ministry for Education and Research (BMBF) in the FitMultiCell project (grant number 031L0159A), the German Research Foundation (DFG) under Germany’s Excellence Strategy EXC 2151 - 390873048 (ImmunoSensation2) and EXC 2047 - 390685813 (Hausdorff Center for Mathematics), and a Schlegel Professorship by University of Bonn. SK is funded by the ReinfChemo project (grant number 031L0270). FF is supported by the Francis Crick Institute, which receives its core funding from Cancer Research UK (CC2242), the UK Medical Research Council (CC2242), and the Wellcome Trust (CC2242).

## Author contributions

DW, FF, PS, YS, MH, SK, and JH together developed the MATLAB toolbox. DW did the validation, checked the computational accuracy of the toolbox, and did the optimization and results analysis with the help of PS, SK, and JH. SK further evaluated experimental data from a previous study and developed the extrinsic apoptosis model. JH, SK and RE supervised the work and obtained funding for this study. DW, SK and JH wrote the first draft of the manuscript. All author read and approved the final manuscript.

## STAR METHODS

- KEY RESOURCES TABLES
- RESOURCE AVAILABILITY
  – Lead Contact
  – Materials Availability
  – Data and Code Availability
- METHOD DETAILS
  – Joint Likelihood and gradient
  – Gradient of the joint likelihood
  – Conversion reaction model
  – Extrinsic apoptosis model
  – Experimental procedures, data processing, and observable definitions
  – Single-cell time-lapse data
  – Single-cell snapshot data
  – Derivation of penalty terms based on the assumption of normally distributed random effects

## METHOD DETAILS

### Mathematical model

In this study, we describe heterogeneous cell populations using nonlinear mixed-effects models. The time evolution of the concentration of biochemical species in each individual cell is modelled using a system of ordinary differential equations (ODEs),

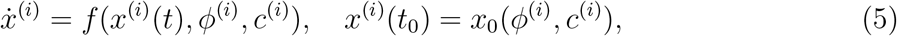

with time *t*, state 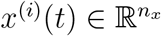, parameters 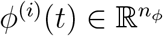, and covariates *c*^(*i*)^. The vector field *f* describes the time evolution of the cell state and is assumed to be Lipschitz continuous. The initial condition is defined via the function *x*_0_, which allows for parameter dependence and constraints such as stationarity. Decision processes of individual cells are modelled using trigger functions *g*(*x*^(*i*)^(*t*), *ϕ*^(*i*)^, *c*^(*i*)^). An event occurs if the corresponding trigger function crosses zero (see section on *Single-cell time-to-event data*.

Cells differ with respect to covariates *c*^(*i*)^ and parameters *ϕ*^(*i*)^. The covariates capture known differences, e.g., in the experimental conditions. The parameters capture unknown differences and are modelled as a linear combination of the fixed effects *β* and the random effects *b*,

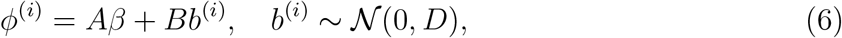

with covariance matrix 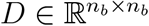, and design matrices 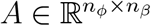 and 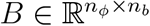, where *n*_*ϕ*_, *n*_*β*_ and *n*_*b*_ are the dimensions of the mixed effects, the fixed effects and the random effects. The design matrices *A* and *B* are often identity matrices or rectangular matrices with 0 or 1 entries.

In this study, we consider only problems with normally distributed random effects. Yet, this is not particularly restrictive as the nonlinearity of the vector field *f* allows for transformations of the normally distributed random variables. For biochemical reaction networks, kinetic rates are often assumed to be log-normally distributed. This can be achieved by using exp(*ϕ*^(*i*)^) in *f* (see, e.g., (Fröhlich et al., 2018a)).

Experiments might not provide direct access to the cell state *x*^(*i*)^(*t*). Therefore, we consider observation functions *h*,

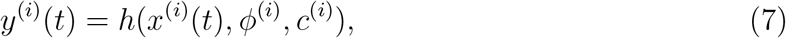

with 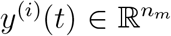 denoting the vector of observables with dimension *n*_*m*_. Furthermore, we consider time-to-event data, which provide the time point *z*^(*i*)^ where the value of a trigger function *g*(*x*^(*i*)^, *ϕ*^(*i*)^, *c*^(*i*)^) crosses zero (Fröhlich et al., 2016). We note that the observation function and even the dimensionality of the observable and the can depend on the experiments. Furthermore, an event might occur or not occur depending on the experimental condition and the cell parameters.

Nonlinear mixed-effect models provide descriptions at both the cell and population levels. At the cell level, the parameters are the mixed effects, 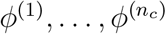, where *n*_*c*_ is the number of cells, and drawn from a probability distribution. At the population level, the parameters are the fixed effect *β* and the covariance matrix of the random effect *D*, which might be parameterized by a vector *δ* such that the mapping *δ* ↦ *D*(*δ*) ensures positive definiteness of *D*.

### Mathematical formulation of likelihood functions

To estimate the parameters of nonlinear mixed-effect models of heterogeneous cell populations from single-cell time-lapse data, population average data, and snapshot data, we formulate in the following the individual likelihood functions as well as the joint likelihood function. Furthermore, we outline numerical schemes to evaluate or approximate the likelihood functions.

To simplify the notation, we skip the dependence on the covariates *c*^(*i*)^ and consider only a single experimental condition. The generalization is discussed below and covered by our implementation.

#### Likelihood function for single-cell time-lapse and time-to-event data

We consider single-cell time-lapse and time-to-event data:

##### Single-cell time-lapse data

consist of measurements of the observables *y*_*k*_, *k* = 1, …, *n*_*m*_, for the time points *t*_*j*_, *j* = 1, …, *n*_*t*_. Assuming independent normally distributed measurement noise, the measured values are given by

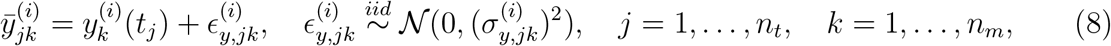

with 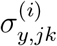 denoting the standard deviation of the measurement noise. The likelihood function for observing these measured values for a given single-cell parameter *ϕ*^(*i*)^ is

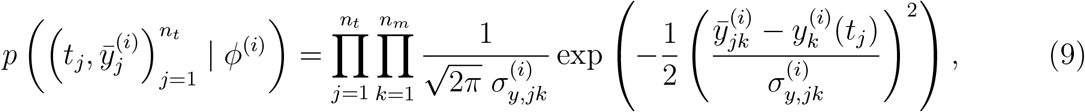

with measurement 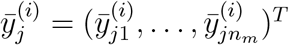, and parameter-dependent model simulation 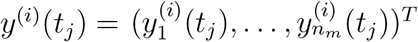.

##### Single-cell time-to-event data

provide information about the time point at which a trigger function *g*(*x*^(*i*)^(*t*), *ϕ*^(*i*)^, *c*^(*i*)^) crosses zero. Yet, as the recording is subject to uncertainty, so is the measured time point 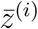. Assuming that the noise is normally distributed, the measured event time is given by

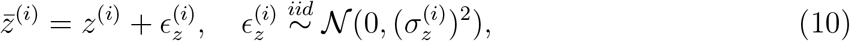

where *z*^(*i*)^ is the root of the trigger function *g* and thus depends on states, parameters, and covariates. 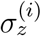 denotes the standard deviation of the measurement noise for the event observation, which can be individual specific. If the event is not recorded, this means that the threshold was not reached before the end of the observation. Here, we write the likelihood function of observing these measured event time for a given single-cell parameter *ϕ*^(*i*)^ as

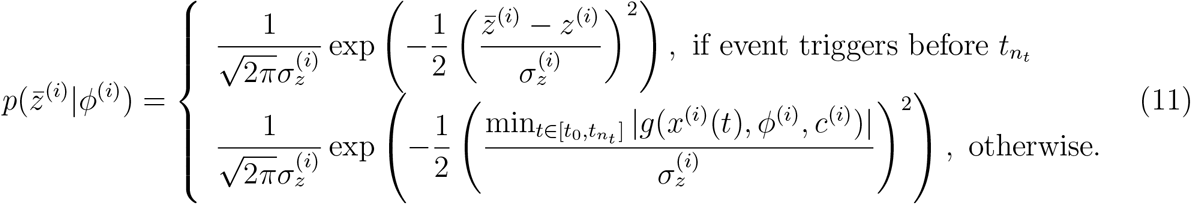

The second case provides a penalty term for the case that in the experiment an event has been observed but the simulation does not capture it. For details, we refer to (Fröhlich et al., 2016).

The likelihood of the single-cell datasets consisting of time-lapse and time-to-event data,

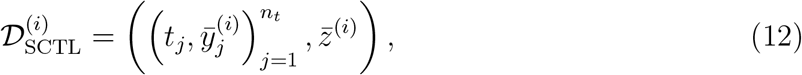

and can for a given single-cell parameter *ϕ*^(*i*)^ be written as the product of the likelihoods of the individual data types:

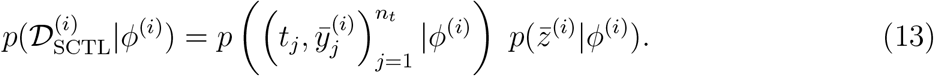

Yet, as the single-cell parameter *ϕ*^(*i*)^ is unknown and not observed, we have to marginalize over *ϕ*^(*i*)^ to assess how likely the single-cell datasets are for a given population with parameters *β* and *D*. The individual marginal likelihood of the single-cell data 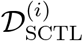 for given population parameters *β* and *D* is

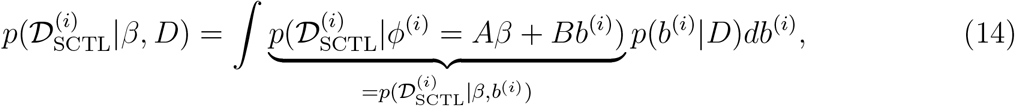

with *p*(*b*^(*i*)^ | *D*) being the multivariate normal density and the integral goes over the multidimensional support of the random effects. As the integrant is a function that depends on the solution of an ODE, it is in most cases not analytically tractable. Hence, the *n*_*b*_-dimension integral has to be approximated numerically, which we discuss further in subsequent paragraphs.

The overall marginal likelihood for a collection of single-cells, assuming independence of the measurements for individual cells, is characterized by the data

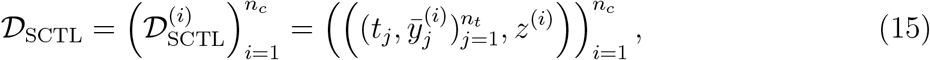

and given by

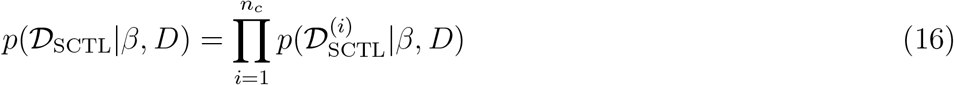

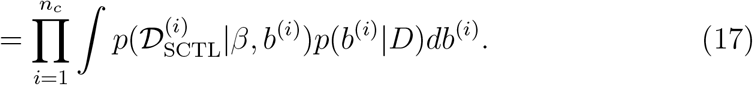

Hence, *n*_*c*_ integrals need to be computed and multiplied. The product can be numerically unstable, in particular for *n*_*c*_ ≫ 1. Therefore, we will usually work with the negative log-likelihood function, turning products into sums

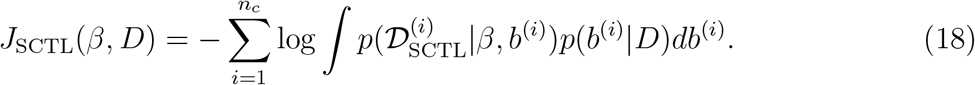

The evaluation of the negative log-likelihood function, becomes increasingly challenging as the number of cells *n*_*c*_ and the number of random effects *n*_*b*_ increases. In particular for *n*_*b*_ ≥ 4, there is only a limited number of suitable and computationally efficient integration routines.

#### Likelihood function for single-cell snapshot data

We consider single-cell snapshot data that provide noise-corrupted measurements of individual cells (8). In contrast to single-cell time-lapse data, individual cells are only observed once, yielding a dataset

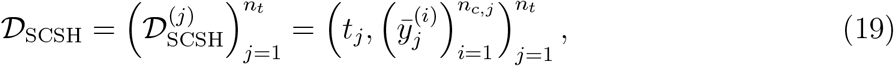

with 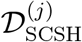 denoting the single-cell snapshot data for time point *t*_*j*_ consisting of *n*_*c,j*_ observations of individual cells.

In principle, we could use for the single-cell snapshot data a similar likelihood formulation as for the single-cell time-lapse data. Yet, as the number of cells observed in single-cell snapshot experiments is much larger, the computational complexity for evaluating this likelihood would be massive.

In this study, we formulate the parameter estimation problem for single-cell snapshot data in terms of their statistical moments. The statistical moments, in particular mean and variance, are descriptive statistics of the state of the cell population. The measured mean and variance of the *k*-th observable for a snapshot 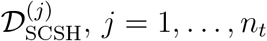 are given by:

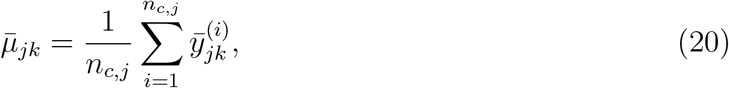

and

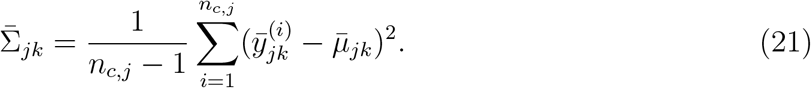

These measured quantities need to be compared to the respective model predictions, which can be obtained by marginalizing over the latent parameters and the measurement noise. The time- and parameter-dependent model expression for the mean is

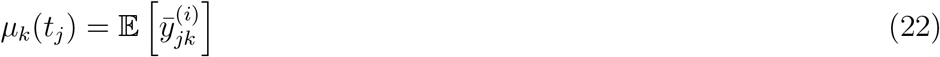

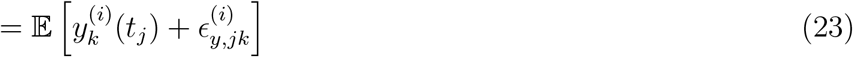

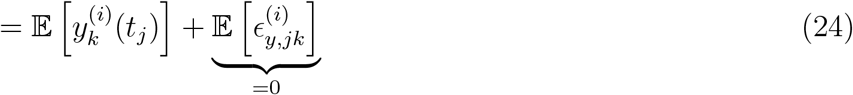

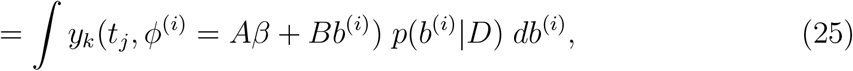

with the second line following from the definition of the single-cell measurement (8), the third line using linearity of the expectation operator and that the mean of the measurement noise is zero, and the last line using the expression of the expected value in terms of an integral over the random effects. The variance is

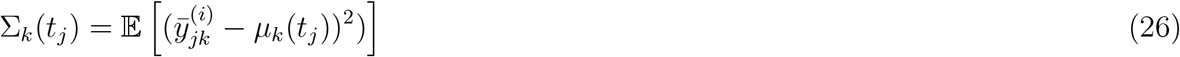

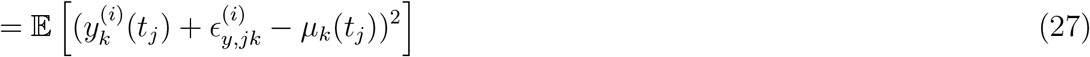

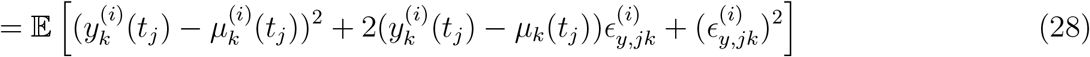

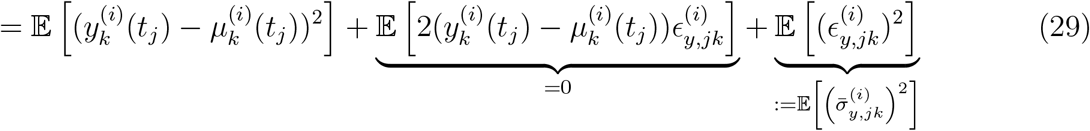

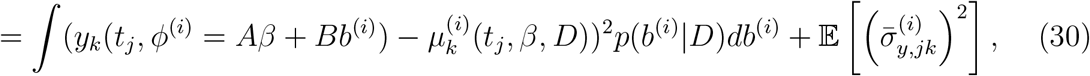

with the second line following from the definition of the single cell measurement (8), the third line providing a reformulation, the fourth line using independence of the random variables and the variance of the measurement noise, and the last line using the expression of the expected value in terms of an integral over the random effects. Note that population variance is not simply the variance of the observable across the population but inflated with the variance of the measurement noise. The variance of the measurement noise, 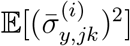, is defined by the corresponding noise distribution and can be computed also by integration over the noise variances for different random effects. If the noise variance is independent of the random effect, 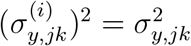, it holds that 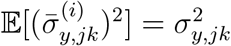.

The likelihood of observing sample means, 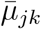, and sample variance, 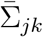, is determined by the sample sizes *n*_*c,j*_ as well as noise on the level of the experimental parameters. While the former goes to zeros as the sample sizes *n*_*c,j*_ increase, noise resulting from differences between experiments (e.g., in sample handling, batches of reagents, temperature, etc.) is independent of the sample sizes. Our assessments suggest that given the large number of cells considered in our experiments, the sampling error is mostly negligible, such that the experimental error is the key factor. We assume here that this experimental error is additive,

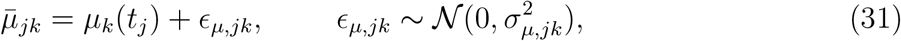

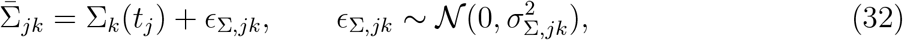

with noise standard deviations *σ*_*µ,jk*_ and *σ*_Σ,*jk*_. Using this noise model, the likelihood function for SCSH data based on the statistical moments is

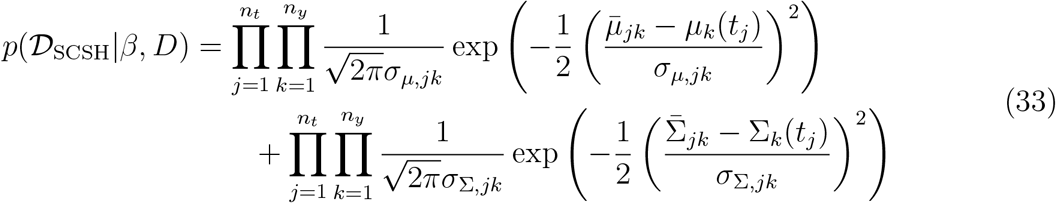

with *µ*_*k*_(*t*_*j*_) and Σ_*k*_(*t*_*j*_) depending on the population parameters *β* and *D*. The likelihood can be extended to co-variances and higher-order statistical moments. Yet, our analysis of alternative population models shows that the information content quickly saturates (Kazeroonian et al., 2013).

The numerically robust evaluation of the likelihood functions for the time-lapse and the time-to-event data becomes, due to the products, increasingly challenging as the number of time points, *n*_*t*_, and the number of observables, *n*_*y*_, increase. Therefore, we consider in the following the negative log-likelihood function

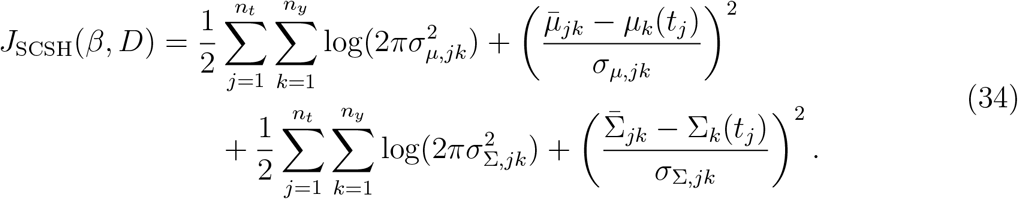

The summation decreases the risk of overflow and underflow during the numerical evaluation compared to the multiplication.

The evaluation of the likelihood function, respectively, the negative log-likelihood function, of single-cell snapshot data requires only the evaluation of the mean and variance of the cell population. Hence, the calculations are independent of the number of observed cells, *n*_*c,j*_, and even measurements with tens of thousands of cells can be easily considered.

#### Likelihood function for population-average data

We consider population-average data, which provide noise-corrupted measurements of the average abundance of a biochemical species within the cell population. The data provide an average over a finite number of cells, however, the number of cells is usually in the thousands, meaning that the sampling error can be disregarded and – as for the statistical moments of the single-cell snapshot data – the noise on the level of the experimental parameters is more critical. Assuming independent normally distributed measurement noise for the overall measurement process, the measured population averages are given by

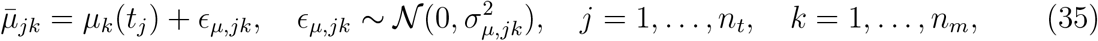

and the dataset is denoted by

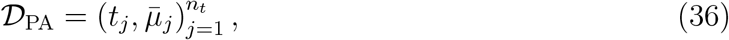

with 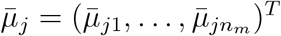.

The likelihood function for population-average data is given by

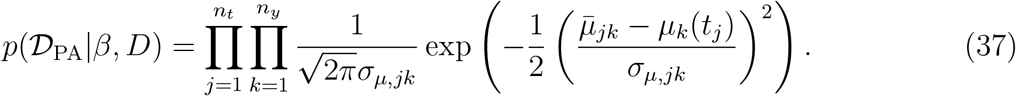

This is similar to the likelihood function for single-cell snapshot data but lacks the variance information. Accordingly, the negative log-likelihood function is given by

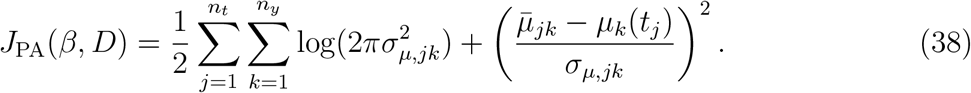

Note here that the standard deviation of the measurement noise can differ between single-cell snapshot and population average data, although this is not indicated by the notation.

Remark: Population-average data can differ from single-cell snapshot data in several important ways:

- **Unequal cell contributions:** Individual cells may contribute differently to the population-level measurement. For example, if the experimental technique measures the total abundance of a biochemical species (in contrast to a single-cell concentration), and cell volumes vary, it is essential to perform calculations on the level of molecule numbers per cell rather than concentrations.
- **Unknown sample size:** The number of cells contributing to the population-average measurement may be unknown, preventing direct normalization. In such cases, an additional scaling factor must be estimated during model fitting, or a proper normalization needs to be performed.

#### Joint likelihood function

In this study, we employ nonlinear mixed-effects models to characterize heterogeneous cell populations by integrating different data types. Specifically, we combine single-cell time-lapse, single-cell snapshot, and population-average datasets. Assuming (i) independent measurements across cells and (ii) independent noise, the joint likelihood is the product of the data-type specific likelihoods,

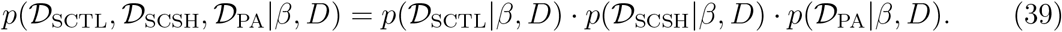

Accordingly, the negative log-likelihood for the combined dataset is given by the sum of the negative log-likelihoods of the individual datasets,

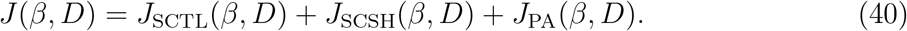

This joint likelihood and negative log-likelihood function readily extends to multiple experimental conditions and alternative observation functions: likelihoods from different conditions are multiplied, and each observation model can be adapted within its respective likelihood term. Moreover, the noise distribution can be flexibly specified—from additive normal to multiplicative log-normal or Laplace—provided independence assumptions hold (Maier et al., 2017).

### Mathematical formulation of parameter estimation problem

The quantitative description of heterogeneous cell populations requires knowledge of the population parameters, namely, the fixed effects *β* and the covariance matrix of the random effects *D*. As many of these parameters cannot be accessed experimentally, we employ maximum likelihood estimation to assess them. The maximum likelihood estimates of the population parameters are obtained by maximizing the joint likelihood for the different datasets

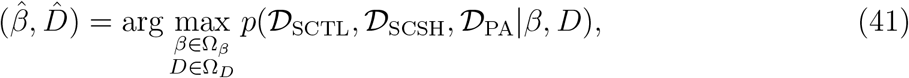

or by minimizing the negative log-likelihood

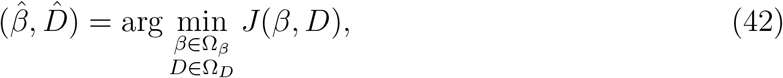

with Ω_*β*_ and Ω_*D*_ indicating the sets of admissible parameter values.

A key challenge for optimization methods are equality and inequality constraints. Therefore, reparameterizations are used to ensure symmetry, *D* = *D*^*T*^, and positive definiteness, *D* ≻ 0, of the covariance matrix:

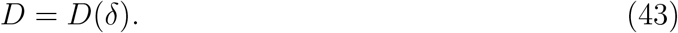

Common choices include the Cholesky decomposition, the spherical parameterization, and the Givens parameterization. In this study, we employ a proposed parameterization based on Lie algebra (Adlung et al., 2021).

The collection of estimated population parameters will be denoted by *θ*, with

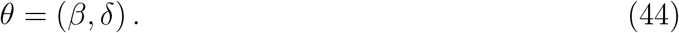

Following this notation, the reparametrized maximum likelihood estimation problem can be stated as

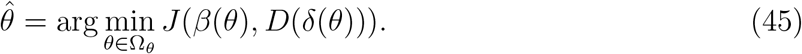

The admissible set of *θ*, Ω_*θ*_, is constructed from the admissible sets of *β* and *δ*. From the maximum likelihood estimate 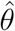 the maximum likelihood estimates for fixed effects, 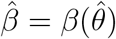, the covariance matrix parametrization, 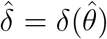, and the covariance matrix, 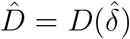, can be computed.

In this study, we solve the nonlinear and non-convex optimization problem (45) using multi-start local optimization. The local optimization is conducted using gradient-based methods as this has been shown to be computationally efficient for many application problems in systems biology (Villaverde et al., 2019). Implementation details are provided below.

We note that standard estimation methods for nonlinear mixed-effect models such as the stochastic approximation expectation-maximization (SAEM) algorithm are not directly applicable for the case of multiple different data types. If merely single-cell time-lapse data were considered, SAEM algorithms would be applicable.

### Numerical approximation of likelihood functions

To determine the maximum likelihood estimate 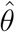, the negative log-likelihood functions need to be evaluated. As the expressions contain high-dimensional integrals, we use tailored numerical approximation schemes.

In the following, we present the respective numerical description and the relevant implementation details. We note that numerical simulations and sensitivity computations are performed using the SUNDIALS package CVODES (Serban and Hindmarsh, 2005).

#### Numerical likelihood approximation for single-cell time-lapse and time-to-event data

The negative log-likelihood function of single-cell time-lapse and time-to-event data is the sum of the negative log-likelihood functions for individual cells, 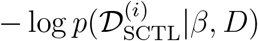, As the single-cell parameters are unknown, these likelihood functions for individual cells are defined as the integral of 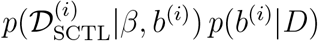 over the random effect *b*^(*i*)^.

To approximate the *n*_*c*_ integrals—one for each cell—we use the Laplace approximation. The Laplace approximation is well established for nonlinear mixed-effects models (Tierney and Kadane, 1986) and its construction follows a two-step procedure:

- Step 1: an estimate for the cell-specific random effect, *b*^(*i*)^, is obtained by maximizing 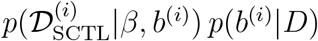 for given values of *β* and *D*. The objective function is the product of the likelihood of the single-cell data and the conditional probability of the random effect. Indeed, this is the integrand of the integral that we aim to approximate. For numerical robustness, we minimize the negative log-transform the objective function,

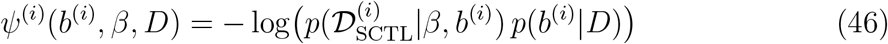

and solve the equivalent problem:

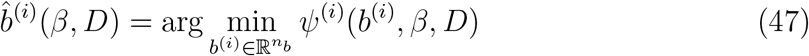

for the current values of *β* and *D*. The estimate 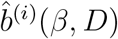 can be interpreted as the Maximum A Posteriori (MAP) estimate of the single-cell parameters given *β* and *D*.
- Step 2: a second-order Taylor series expansion of the negative log-transformed integrand, *ψ*^(*i*)^(*b*^(*i*)^, *β, D*), at the estimate 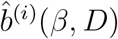. This approximation is given by

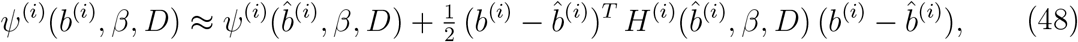

with

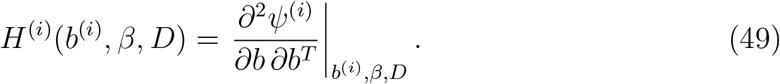

The gradient 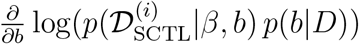 is zero at the estimate 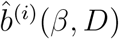 since it is the mode of the function *ψ*^(*i*)^(*b*^(*i*)^, *β, D*), such that the first order term vanishes. Furthermore, the Hessian *H*^(*i*)^(*β, D*) is, due to the term 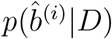, essentially always positive definite (even if the ODE model is non-identifiable).
- Step 3: an approximation of the integral is obtained by integrating the Taylor series expansion of the integrand. As this Gaussian integral can be solved analytically, we obtain

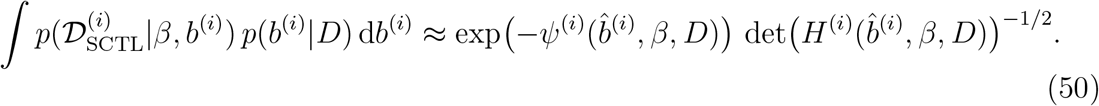

The approximation of the likelihood for a collection of cells is given by the product of the approximations for the individual cells,

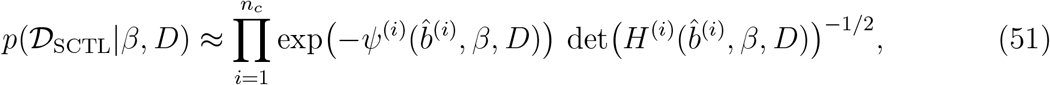

and the corresponding, approximated negative log-likelihood function is

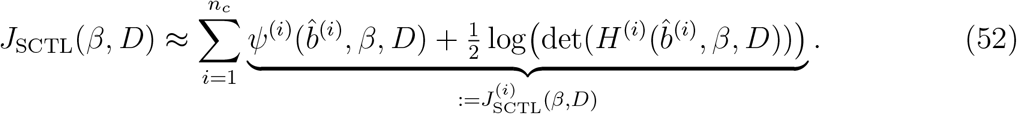

For the numerical evaluation of the approximated negative log-likelihood, the inner optimization problems have to be solved and the Hessian (49) has to be evaluated. Here, we solve the optimization problem using gradient-based multi-start local optimization. The starts are initialized at (1) the optimal point of the previous iteration of the outer optimization and (2) random points sampled from *p*(*b* | *D*). In total, 10 starts are performed. For the optimal point, we compute the Hessian using forward sensitivity equations and the implicit function theorem (see (Fröhlich et al., 2016)).

#### Numerical likelihood approximation for single-cell snapshot data

The negative log-likelihood function of the single-cell snapshot data depends on the population means and variances, *µ*(*t*) and Σ(*t*). As these expressions are for most models not accessible in closed form, we compute approximations using Monte Carlo sampling. We denote the sampling-based estimates by

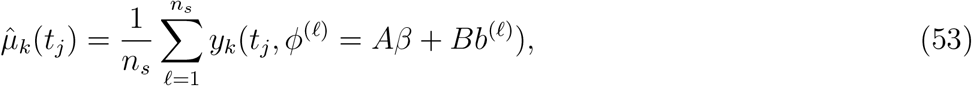

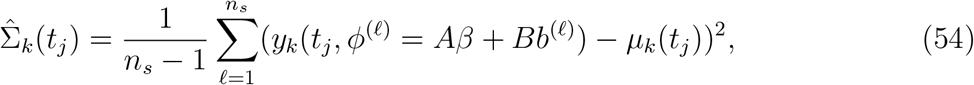

with sample size *n*_*s*_, ODE solution *y*_*k*_(*t*_*j*_, *ϕ*^(*ℓ*)^ = *Aβ* +*Bb*^(*ℓ*)^), and random effect *b*^(*ℓ*)^ ∼ 𝒩(0, *D*). The sampling-based estimates depend on the population parameters *β* and *D*. To ensure differentiability with respect to *β* and *D*, instead of re-sampling the random effects in each iteration of the optimization process, we employ a two step construction of the Monte Carlo approximation:

- Step 1: Standard normally distributed random vectors *r*^(*ℓ*)^ are sampled before the estimation run,

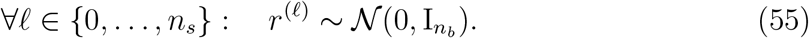

These values are stored and remain constant for the course of an individual optimization.
- Step 2: In each optimizer iteration, the random effects are computed by multiplying the fixed, standard normally distributed random vectors with the matrix square root of the covariance matrix of the random effects, *D*,

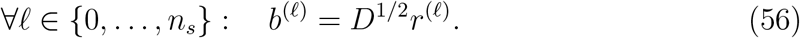

This yields in each iteration random effects which are distributed according to 𝒩(0, *D*). The matrix *D*^1*/*2^ is computed using the Cholesky decomposition.

Given the random effect *b*^(*ℓ*)^, the ODE model can be simulated to obtain the state trajectories and outputs *y*_*k*_(*t*_*j*_, *ϕ*^(*ℓ*)^ = *Aβ* + *Bb*^(*ℓ*)^), from which an approximation of the mean and the variance can be derived.

As an alternative to the Monte Carlo approximation, we also considered the sigma-point approximation. Sigma-point methods provide deterministic quadrature rules for approximating the moments of transformed random variables (Julier, 2002), particularly when the transformation is nonlinear–as is the case when propagating random effects through an ODE model and observation function.

In the sigma-point approximation, a fixed number of carefully chosen evaluation points (so-called sigma points) are selected based on the distribution of the random effects. These sigma points are propagated through the model to estimate the first and second moments:

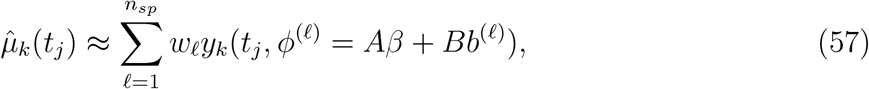

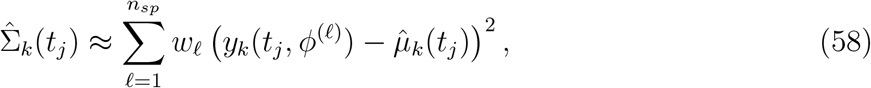

where *b*^(*ℓ*)^ and *w*^(*ℓ*)^ are the sigma points and associated weights, respectively, and *n*_*sp*_ denotes the number of sigma points. We employ the unscented transform to generate the sigma points and weights, using *n*_*sp*_ = 2*n*_*b*_ + 1 (Wang et al., 2019). This method is designed to exactly reproduce the mean and covariance up to second order for Gaussian-distributed inputs. The sigma points are generated deterministically from the mean and covariance of the random effects, which enables fully differentiable approximations without sampling noise.

Compared to Monte Carlo methods, sigma-point approximations can achieve high accuracy with fewer model evaluations, especially in low-dimensional settings. However, for the consider application we found the accuracy to be insufficient and rather employed Monte Carlo approximations.

#### Numerical likelihood approximation for population average data

The negative log-likelihood function for population-average data depends on the population means, *µ*(*t*). Since closed-form expressions for *µ*(*t*) are generally not available for most models, we approximate it using the same Monte Carlo sampling strategy employed for single-cell snapshot data.

#### Numerical likelihood approximation for joint likelihood function

The joint likelihood function is approximated by multiplying the likelihood contributions of the individual data types. Correspondingly, the joint negative log-likelihood function is approximated by summing the individual negative log-likelihoods.

As each of the individual likelihood functions is itself an approximation, the joint likelihood function is also subject to approximation error. However, all approximations improve as datasets become more informative and as inter-individual variability decreases. In particular, the Laplace approximation becomes more accurate when the posterior distributions of the random effects are sharply peaked.

It is important to note that the observation function *h* may differ between data types. Consequently, simulations used for one data type cannot always be reused for another. Separate simulations may be required for each likelihood component to ensure consistency with the corresponding observation models.

### Gradient computation for likelihood functions

To ensure numerical efficiency and robustness during parameter estimation, we compute gradients of the (approximated) negative log-likelihood function using forward sensitivity analysis in combination with the implicit function theorem. This approach avoids the numerical instabilities and computational overhead associated with finite difference schemes, which are particularly problematic for high-dimensional or stiff systems.

The computation of the gradient with respect to the population parameters *θ* = (*β, δ*) requires careful treatment of several model characteristics:

- The model exhibits a hierarchical structure with complex multi-level dependencies. Accordingly, the chain rule must be applied systematically to derive expressions for the gradient. A precise distinction between total and partial derivatives is required; otherwise, the resulting gradients may be incorrect. Even small inaccuracies can substantially impair optimization performance.
- For all data types, the model outputs depend on the solutions of ODE systems. Thus, derivatives with respect to parameters are computed using forward sensitivity equations, which efficiently propagate parameter perturbations through the dynamics.
- For single-cell time-lapse and time-to-event data, the approximated likelihood is constructed via the Laplace approximation, which depends on the solution of an inner optimization problem for the random effects. As a result, the gradient includes contributions from the optima 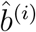 of these inner problems, necessitating the use of the implicit function theorem to correctly differentiate through the optimization result.

The full mathematical expressions for the gradients are complex and span multiple pages. Rather than reproducing them here, we refer the reader to our accompanying implementation, which includes symbolic expressions and implementation notes. These materials clarify how gradients are composed from partial derivatives of model outputs, observation functions, and likelihood terms.

This modular and analytically grounded gradient computation underpins the gradient-based optimization algorithms used in our study and ensures scalable performance across diverse model structures and data configurations.

### Model and synthetic data for conversion process

To evaluate the proposed methods under controlled conditions, we considered a simple conversion reaction model, *x*_1_ ⇌ *x*_2_. The model could represent a reversible step in a signal transduction system such as the exchange of a protein between cellular compartments or a reversible conformational change. The dynamics of the system are described by the following system of ordinary differential equations (ODEs) with forward and backward reaction parameters *k*_1_ and *k*_2_:

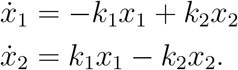

To keep the parameter space low-dimensional, we do not use independent, unknown initial conditions but chose the non-equilibrium conditions *x*_1_(0) = *k*_1_*/*(*k*_1_+*k*_2_) and *x*_2_(0) = *k*_2_*/*(*k*_1_+ *k*_2_). Conceptually, this means that the rates of the process are perturbed at time point *t* = 0.

We use the parameterization *k*_1_ = exp(*ϕ*_1_) and *k*_2_ = exp(*ϕ*_2_), ensuring non-negative reaction rates.

We generated synthetic datasets for each of the three data types considered in this study: *single-cell time-lapse and time-to-event data (SCTL), single-cell snapshot data (SCSH)*, and *population-average data (PA)*. All datasets were simulated using known parameter values, with added Gaussian noise to mimic experimental variability.

Following common experimental assumptions, the PA and SCSH data were assumed to provide *absolute measurements*, whereas the SCTL data were considered to provide only *relative measurements*. To account for this, the SCTL data include an unknown experiment-specific scaling factor, *ϕ*_3_. In addition, time-to-event data were simulated by recording the time 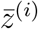 at which the concentration 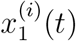 in cell *i* crosses a specified threshold, defined as 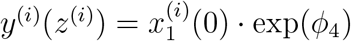.

The detailed configurations for each dataset are as follows:

#### Single-cell time-lapse and time-to-event data

- Number of cells: *n*_*c*_ = 30
- Single-cell measurement: 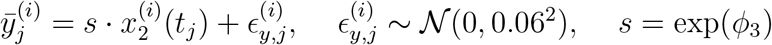
- Time-to-event measurement: 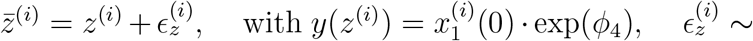 𝒩(0, 0.06^2^)

#### Single-cell snapshot data

- Number of cells: *n*_*c*_ = 100
- Measurement: 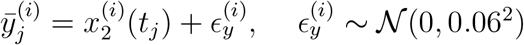
- Measured population mean: 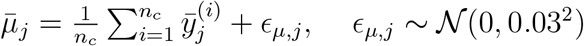
- Measured population variance: 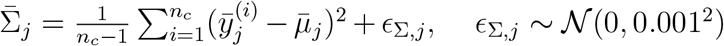

#### Population-average data

- Number of cells: *n*_*c*_ = 100
- Measurement: 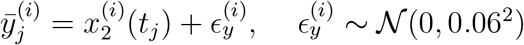
- Measured population mean: 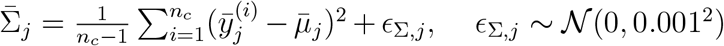

##### Parameter structure

The mixed-effect parameters were defined as:

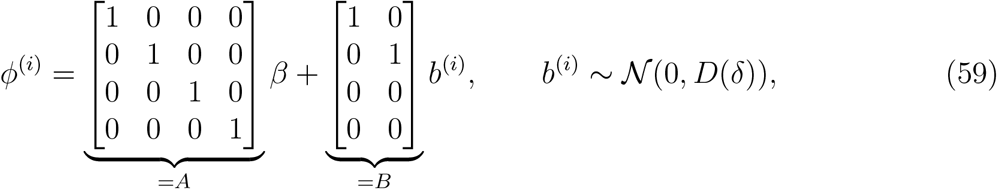

with values

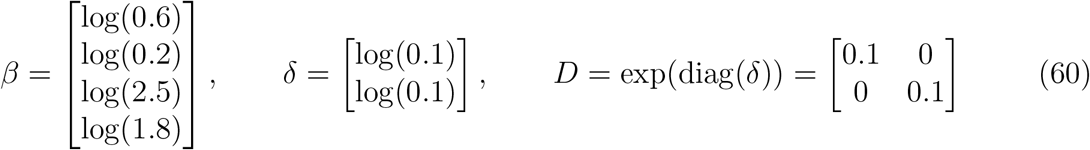

used for simulating synthetic data. This synthetic benchmark provides a controlled environment for method validation, allowing us to analyze parameter recovery and the influence of data type and heterogeneity on inference performance.

### Model and experimental data for extrinsic apoptosis model

As an important application example, we consider a comprehensive model and dataset for extrinsic apoptosis. Extrinsic apoptosis is initiated by extracellular death ligands, such as CD95 ligand (CD95L, also known as Fas ligand) or TRAIL. The binding of CD95L to CD95 (Fas) receptors induces the formation of death inducing signaling complexes (DISCs) (Kischkel et al., 1995). DISCs serve as a platform for the activation of initiator caspases, caspase-8 and caspase-10, that cleave and activate the effector caspases, caspase-3 and caspase-7, and cleave the proapoptotic Bcl-2 family member BID into tBID. Accumulation of tBID induces mitochondrial outer membrane permeabilization (MOMP) that irreversibly triggers activation of effector caspases and cell death (Kallenberger et al., 2014). In type I cells, effector caspase activation by initiator caspases is sufficient for apoptosis, whereas type II cells require indirect effector caspase activation by MOMP (Scaffidi et al., 1998).

By combining experimental data of single-cell caspase-8 activities with population measurements of caspase-8 fragments from immunoblotting experiments, an effective mechanism of caspase-8 activation was determined, in which the cleavage reactions from p55 to p43 and from p30 to p18 are described as interdimeric “trans” reactions, catalyzed by neighbored DISC-bound caspase-8 intermediates, and the reactions from p43 to p18 and from p55 to p30 are intra-dimeric “cis” cleavage reactions, reflected by uni-molecular reaction kinetics (Kallenberger et al., 2014) as illustrated in Fig. 2. The catalytically active fragments p43 and p18 can cleave BID to tBID, which is assumed to induce cell death at a certain threshold concentration. Model equations are given in Supplementary table S1.

In the model, it was assumed that cell-to-cell variability can be exclusively attributed to differences in initial protein concentrations, reflected by a multi-variate log-normal distribution, and that reaction parameters are equal for all cells. Therefore, random effects were included for initial protein concentrations. Additionally, random effects were described for cell volumes and cellular events (apoptosis time points). Reaction parameters were exclusively described by fixed effects.

The model contains in total 24 mixed effect parameter of which 11 are influence by random random effects. The covariance matrix for the random effects is assumed to be diagonal, resulting in 35 population-level parameters. The measurement noise variances of single-cell mean and variance were approximated via experimental replicates.

#### Single-cell time-lapse and time-to-event data

For model fitting, single-cell data were taken or additionally extracted from live-cell imaging data recorded as part of our previous study (Kallenberger et al., 2014). To selectively measure single-cell cleavage activities of the membrane-bound form of caspase-8 (p43) and cytosolic caspase-8 (p18) two cleavage probes, in which a compartment anchor and a fluorescent protein are linked by a caspase-8 specific cleavage site, were developed.

The first probe, Pr_NES_mCherry, contains a nuclear export sequence (NES) and is therefore localized in the cytoplasm, where it can be accessed by p43 and p18. The second probe Pr_ER_mGFP contains a compartment anchor targeting the probe to the endoplasmatic reticulum (ER) and can therefore be accessed by cytosolic p18 only.

A detailed description of the experimental method for quantifying single-cell cleavage activities can be found in (Kallenberger et al., 2014). In short, to measure the cleavage activities of membrane-bound p43 and cytosolic p18, two fluorescent cleavage probes were stably expressed in wild-type HeLa and CD95-HeLa cells. The cleavage probes consisted of a localization sequence for targeting probe molecules to a cellular compartment, the cleavage sequence IETD specifically cleaved by caspase-8, and a fluorescent protein. Upon cleavage, the fluorescent proteins could enter the nucleus (Fig. 2). One of the cleavage probes, Pr_ER_mGFP, contained a truncated version of calnexin to localize it at the endoplasmic reticulum (ER) and make it accessible only for cytosolic p18. The other probe, Pr_NES_mCherry, contained a nuclear export sequence (NES) and was therefore localized in the cytoplasm where it was accessible for p43 as well as p18. The rate equation for the cleaved probe concentration Pr, depending on the concentration of the intact probe PrF and the concentration of caspase-8 was re-arranged to estimate the cleavage activity, resembled by the product of C8 and the rate constant for probe cleavage *k*_cl_,

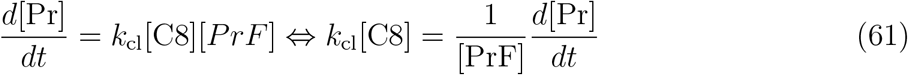

By quantifying the nuclear fluorescence intensities for the cytoplasmic and ER probes (I_NES,ncl_, I_ER,ncl_) and the average intensities in total cell volumes (I_NES,tot_, I_ER,tot_), cleaved probe fractions were determined to infer cleavage activities. After cleavage of probes restricted to the cytoplasm (V_cpl_), fluorescent proteins (mGFP and mCherry) can enter the nucleus (V_ncl_) and equilibrate over the cell (V_c_ = V_cpl_ +V_ncl_). The distribution over the cell results in a dilution by V_c_*/*V_cpl_. We assume that the concentration is related to the measured fluorescence intensity by a scaling factor *c*. Thus, the cleaved probe concentration is related to the nuclear intensity I_ncl_ by

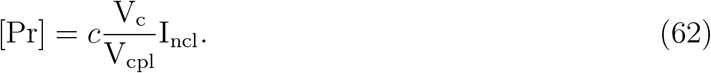

The average cellular intensity is proportional to the sum of uncleaved probe molecules N_PrF_ and cleaved probe molecules N_Pr_ divided by the cellular volume, I_tot_ = *c*(N_PrF_ + N_Pr_)*/*V_c_. The concentration of cleaved probe molecules equals [Pr] = N_Pr_*/*V_cpl_. Therefore, the concentration of uncleaved probe molecules can be estimated by

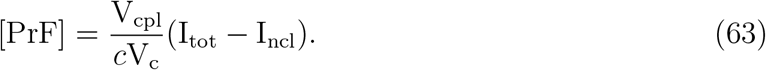

Using (62) and (63), the cleavage activity can be estimated as

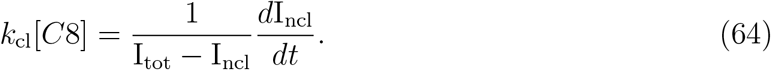

Cleavage of the ER-anchored probe is reflected by cytoplasmic p18,

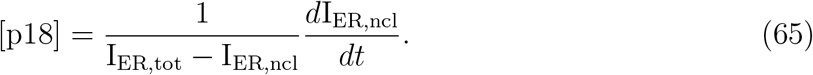

The cytoplasmic probe was cleaved by p43 and p18. Therefore, the cleavage activity of p43 was estimated by subtracting the p18 activity from the cleavage activity indicated by the cytoplasmic probe

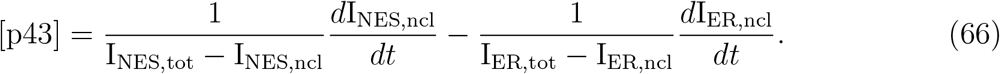

Experimental data sets were recorded in wild-type HeLa cells and in HeLa cells stably over-expressing CD95 (CD95-HeLa). In wild-type HeLa cells, we included 10 single-cell data sets per ligand concentration. Due to the accelerated cell death kinetics resulting from CD95 overexpression, more single-cell data sets could be recorded in CD95-HeLa cells. In these cells, 30 single-cell data sets per ligand concentration were used for model fitting. In all cells, measurements were recorded until apoptosis. These time points defined the cellular events that were used for model fitting. Additionally, cell volumes were determined by confocal microscopy. Cell volumes were used to create a link between experimental magnitudes either proportional to protein concentrations or to absolute amounts of proteins.

Single-cell time-lapse data sets were recorded in wild-type and CD95-HeLa cells at four different CD95L concentrations (wild-type HeLa cells at 500, 2000, 5000 or 10000ng/ml, CD95-HeLa cells at 50, 500, 5000 or 10000ng/l). Standard deviations of the measurement errors of single-cell data were estimated based on fitting smoothing spline functions to recorded time series of fluorescence intensities and error propagation as described in (Kallenberger et al., 2014). The observables are

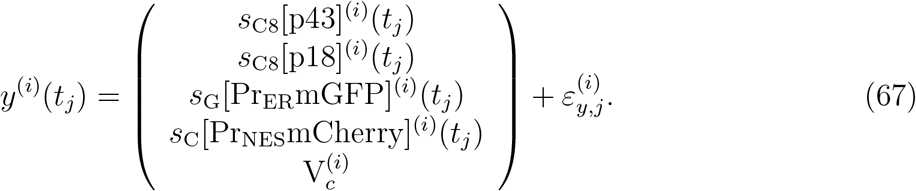

Measurements were collected for individual cells, with the number of time-points being cell-specific. The cell volumes 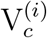 were only quantified at the beginning of the experiment and assumed constant.

#### Population-average data

In this study, we used previously recorded immunoblotting datasets as population average (PA) data (Kallenberger, 2014). For measuring caspase-8 and BID, cell lysates were created from 2.04 · 10^5^ cells per data point. Time series data (p55, p43, p30, p18, BID, tBID) were recorded for CD95-HeLa cells at two ligand concentrations, 50ng/ml and 500ng/ml. Further, by quantitative immunoblotting, average molecule numbers of CD95R, p55, FADD, BID as well as the cleavage probes Pr_NES_mCherry and Pr_ER_mGFP were determined in wild-type HeLa and CD95-HeLa cells. To this end, calibrated protein lysates were used (Kallenberger et al., 2014). Immunoblotting experiments were performed with 3 replicates to estimate standard errors.

In time-resolved experiments, absolute values for immunoblot intensities and the corresponding population averages for each caspase-8 species (p55, p43, p30 and p18) and the two forms of BID (BID and tBID) were normalized by the respective sums of intensities caspase-8,

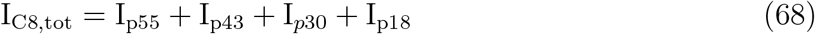

and BID intensities,

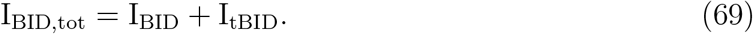

By using immunoblotting band intensity fractions instead of absolute intensities, the introduction of another scaling factor could be avoided. Furthermore, fractions of immunoblotting intensities were used in order to avoid errors from small differences in total protein amounts within immunoblotting samples for different time points that could result from the experimental procedure of sample preparation. Accordingly, measured population averages were related to normalized population averages averages computed from corresponding single-cell variables. To this end, in the model, single-cell concentrations were weighted according to their cell volumes 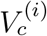. Fractions of caspase-8 or BID species were calculated relative to volume-weighted single-cell concentrations of caspase-8 or BID,

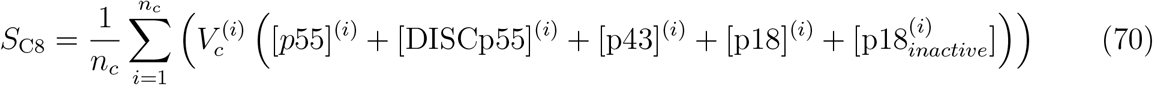

and

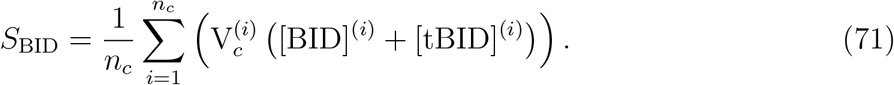

The observables for population averages of caspase-8 species at time points *t*_*j*_ with *j* = 1, …, *n*_*t*_ and errors *ε*_*PA*_ read

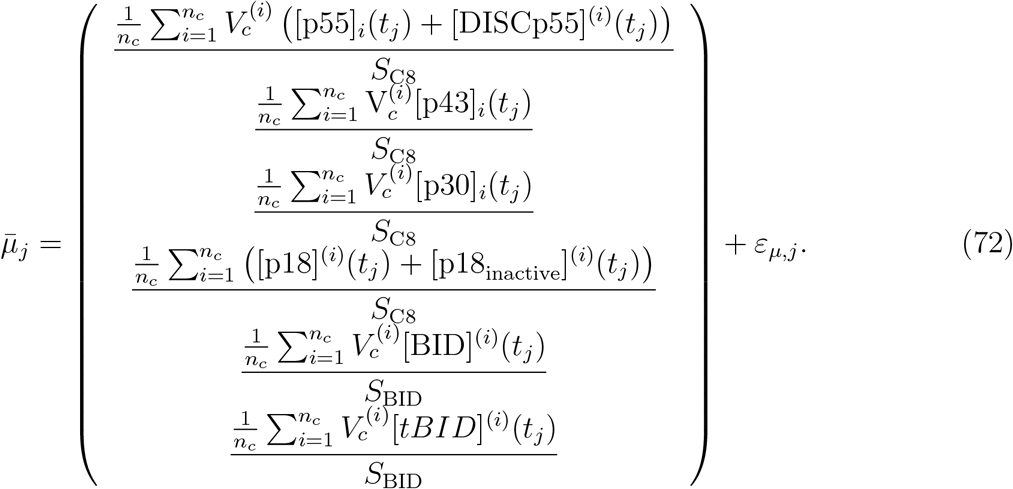

We note that (1) in the population average experiment we do not have access to the measurements for the individual cells but directly observe the indicated sums, and that (2) for the chosen normalization the unknown cell number cancels out.

For p55 and p18, model species that could not be experimentally distinguished (DISC-associated or free p55 as well as active or inactive p18, [p18]^(*i*)^ and [p18_inactive_]^(*i*)^) were collected in single observables. The caspase-8 observables were fitted to ratios of immunoblotting intensities,

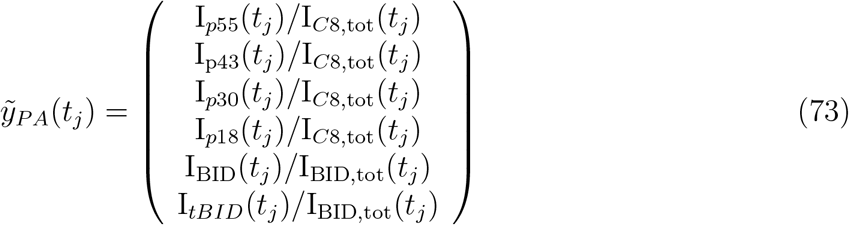

representing fractions of the amounts of caspase-8 or BID species. While the single-cell observable for p18 reflected the corresponding single-cell variable [p18]^(*i*)^ of active p18, the population-average observable derived from the immunoblot band intensity I_p18_ reflected the sum of active as well as inactive p18 and was related to the sum of single-cell variables. Combining single-cell and immunoblotting data facilitated estimating the stability of p18 (Kallenberger, 2014).

#### Single-cell snapshot data

To generate single-cell snapshot (SCSH) data, we re-analyzed previous fluorescence-assisted cell sorting (FACS) experiments (Kallenberger et al., 2014). We estimated variances of initial protein concentrations of CD95R, FADD, p55, BID, Pr_NES_mCherry and Pr_ER_mGFP. FACS experiments were performed for an initial number of 5 · 10^5^ cells per condition. Furthermore, we calculated the variance of cell volumes determined from confocal microscopic image stacks of wild-type and CD95-HeLa cells. To avoid a bias from fluorescent cleavage probes, it was necessary to conduct FACS experiments in wild-type HeLa cells and CD95-HeLa cells without probe expression (Kallenberger et al., 2014). CD95-HeLa were generated from wild-type HeLa cells by stable overexpression of CD95 receptors. Therefore, in the current study, we introduced the simplifying assumption that wild-type and CD95-HeLa cells do not differ in average concentrations of p55, FADD and BID. Different variances, however, were assumed for initial protein concentrations of CD95, Pr_NES_mCherry and Pr_ER_mGFP. FACS experiments were only performed once for p55 and FADD and twice for BID in each cell line. To still estimate the error of quantifying variances *σ*^2^ from FACS experiments, standard deviations were calculated from pairs of measurements of the two cell lines for p55, FADD and BID. The mean of the standard deviations for estimates of initial protein concentration variances was used for weighting residuals in the likelihood function terms for SCSH data.

## Supplementary Tables

**Table S1.**
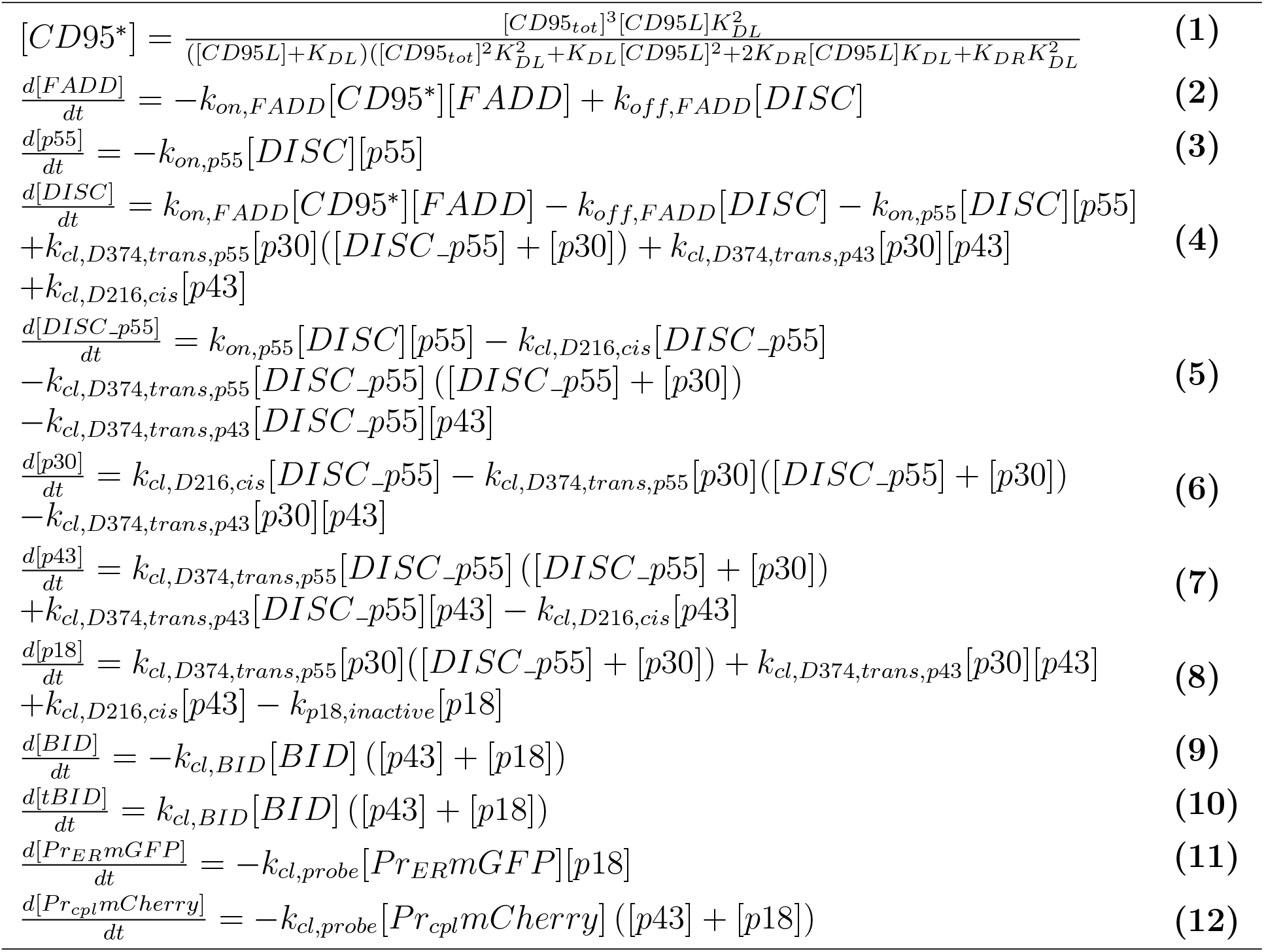
Equations of the extrinsic apoptosis model related to Star Methods.

**Table S2.** Overview of data related to Fig. 5 and Star Methods.

| Dataset | 1 | 2 | 3 | 4 | 5 | 6 | 7 |
| --- | --- | --- | --- | --- | --- | --- | --- |
| Data type | SCTL | SCTL | SCTL | SCTL | SCTL | SCTL | SCTL |
| Cell line | CD95 | CD95 | CD95 | CD95 | wild-type | wild-type | wild-type |
| Cell number | 30 | 30 | 30 | 30 | 10 | 10 | 10 |
| CD95 concentration | 50ng/ml | 500ng/ml | 5000ng/ml | 10000ng/ml | 500ng/ml | 2000ng/ml | 5000ng/ml |
| Dataset | 8 | 9 | 10 | 11 | 12 | 13 | 14 |
| Data type | SCTL | PA | PA | PA, quantitative | PA, quantitative | SCSH | SCSH |
| Cell line | wild-type | CD95 | CD95 | CD95 | wild-type | CD95 | wild-type |
| Cell number | 10 | $2,04 \cdot 10^5$ | $2,04 \cdot 10^5$ | $2,04 \cdot 10^5$ | $2,04 \cdot 10^5$ | $5 \cdot 10^5$ | $5 \cdot 10^5$ |
| CD95 concentration | 10000ng/ml | 50ng/ml | 500ng/ml | 0ng/ml | 0ng/ml | 0ng/ml | 0ng/ml |

**Table S3.**
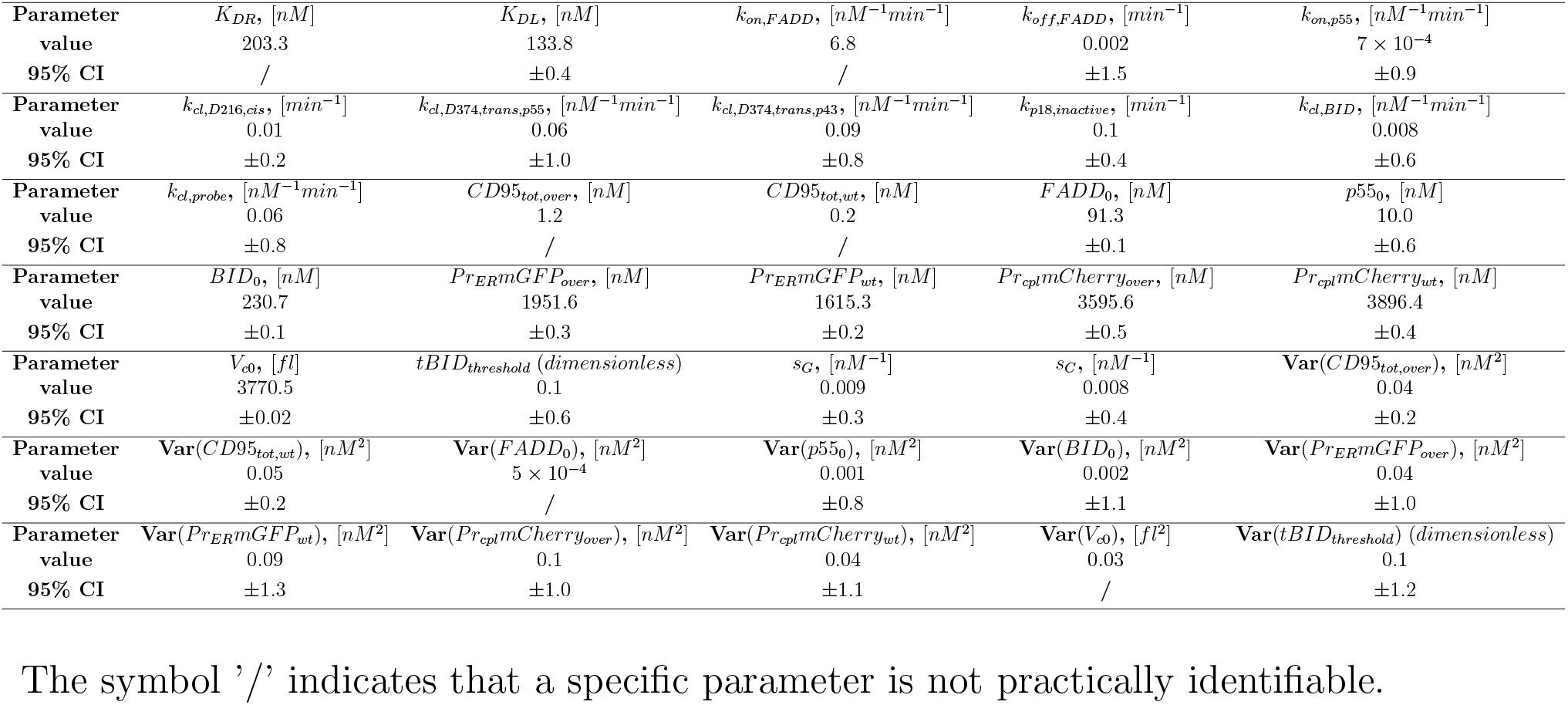
Estimated parameter values and confidence intervals related to Fig. 5 and Star Methods.

## Supplementary Figures

**Supplementary Figure S1:**
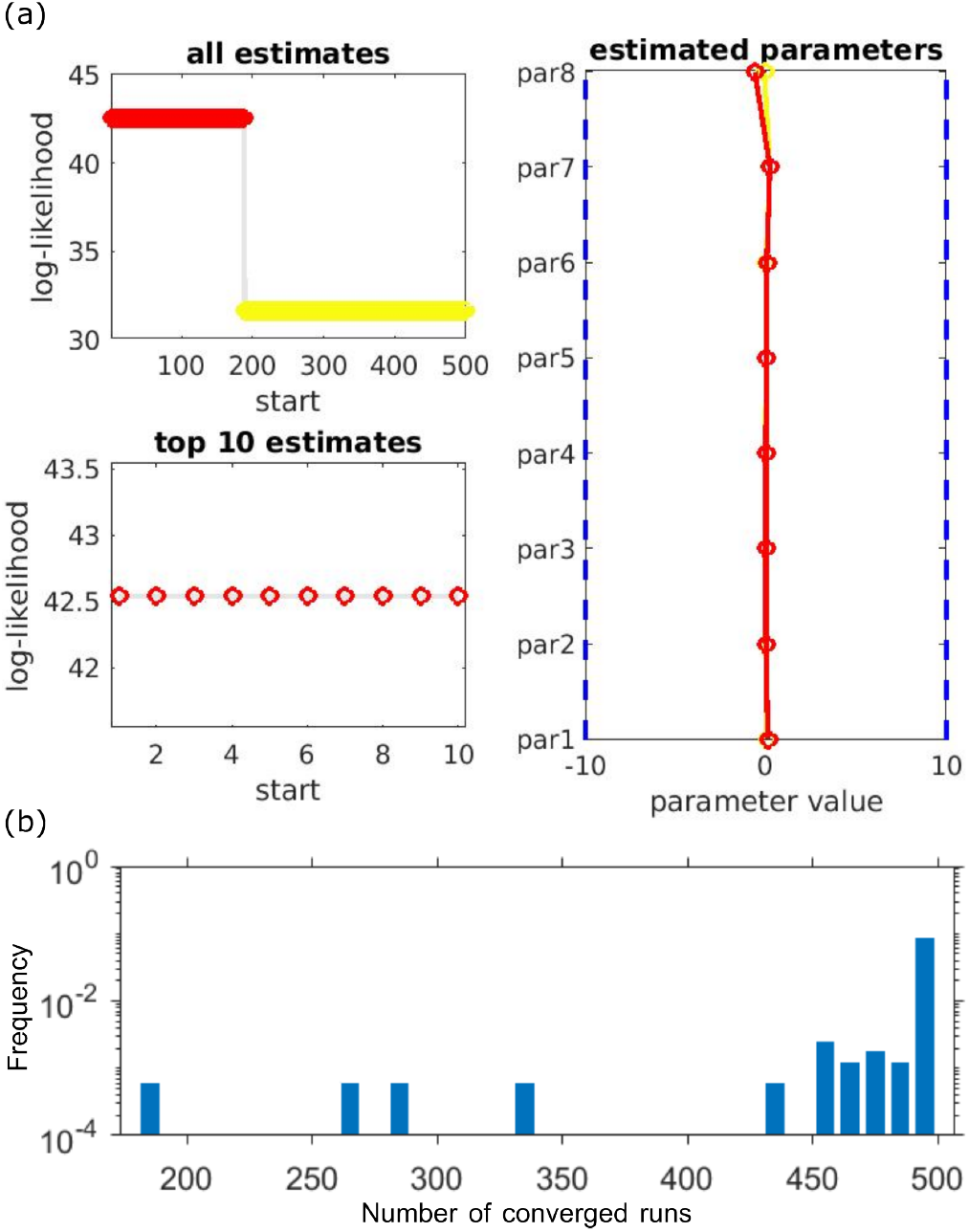
Analysis of robustness of inner optimization results, related to Fig. 2. (a) Left: all/top 10 log-likelihood values across all optimization runs. Right: parameter values across all runs. (b) Frequency of getting a specific number of converged runs when tested on 160 cells with time-lapse data.

**Supplementary Figure S2:**
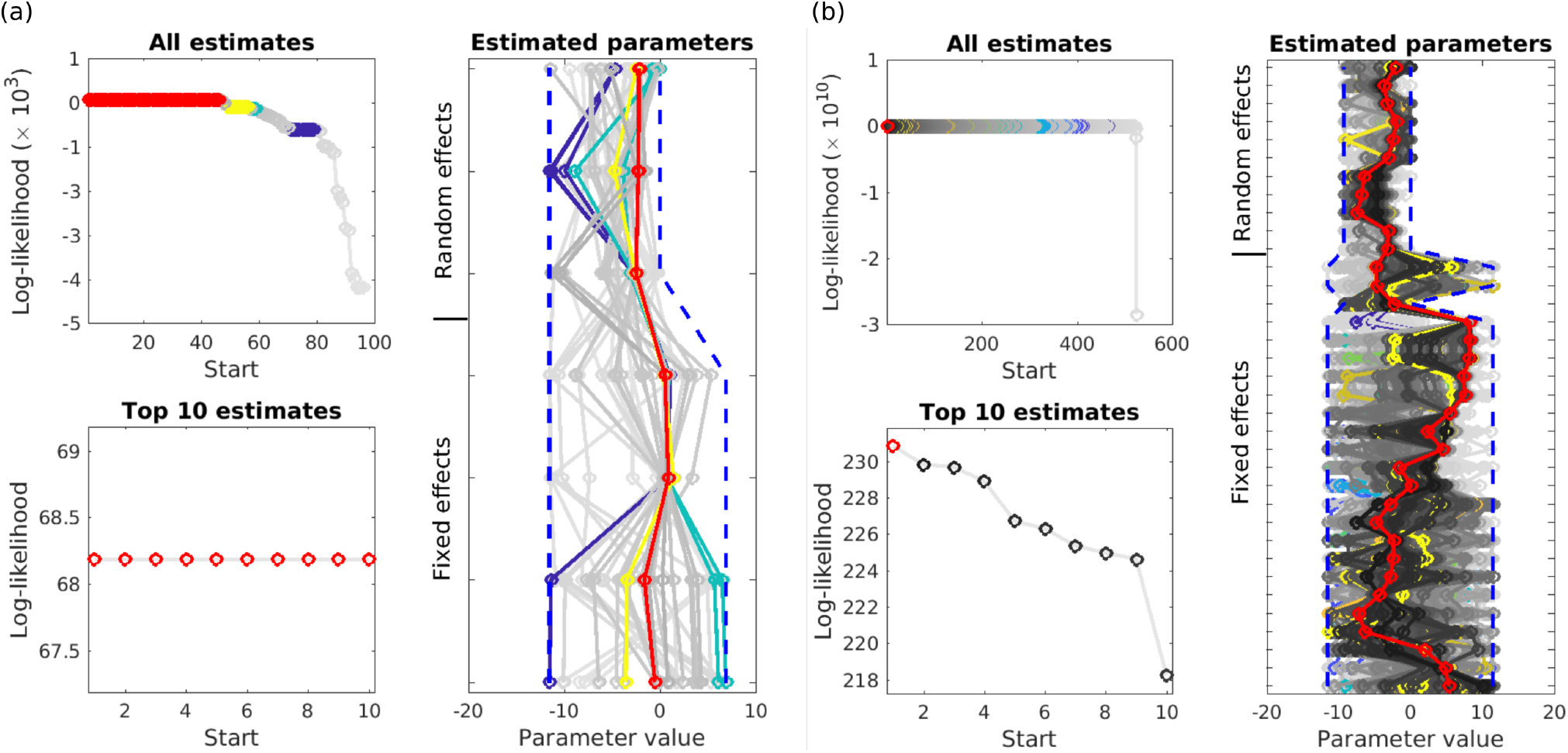
Analysis of reproducibility of optimization results for (a) the conversion reaction model and (b) the apoptosis model, both are for scenario 1. Related to Fig. 5 and Fig. 6. In each subplot, the left two plots show all/top 10 log-likelihood values across all optimization runs. The largest objective function values are colored in red. For other optimization runs, coloring is only applied if multiple optimization runs yielded objective function values within an interval of 0.01 of each other, otherwise a grey color is applied. Failed optimization runs are not shown. Right plot shows the parameter values across all optimization runs.

**Supplementary Figure S3:**
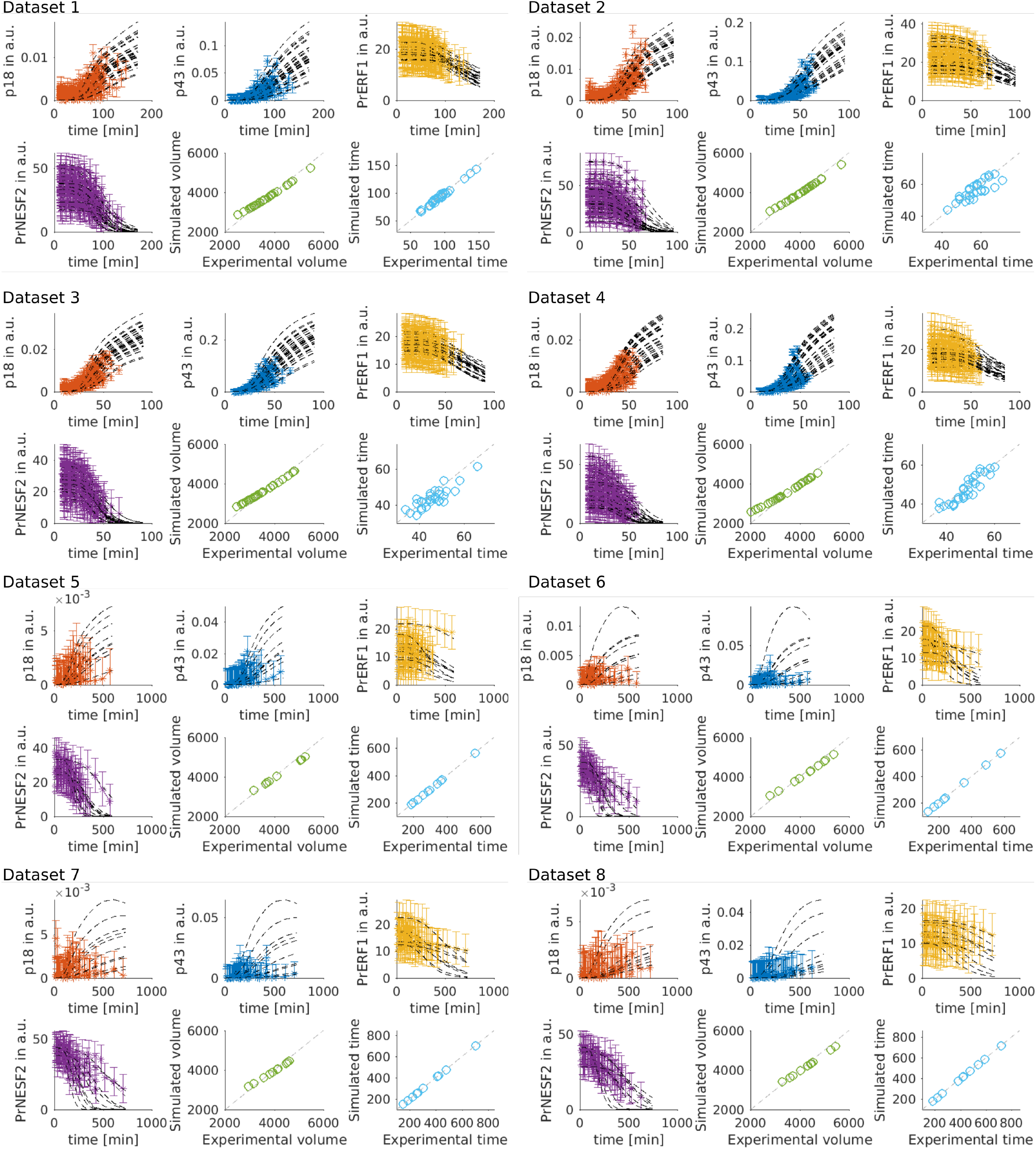
Single cell fittings of the apoptosis model for scenario 1, related to Fig. 5. Different colors are applied for different measurements in one experimental condition. Datasets 1 to 4 comprised a number of 30 CD95-HeLa cells, datasets 5 to 8 a number of 10 wt HeLa cells (see experimental methods for details).

